# Beyond blooms: A novel time series analysis framework predicts seasonal keystone species and sheds light on Arctic pelagic ecosystem stability

**DOI:** 10.1101/2024.03.11.583746

**Authors:** Ellen Oldenburg, Raphael M. Kronberg, Katja Metfies, Matthias Wietz, Wilken-Jon von Appen, Christina Bienhold, Ovidiu Popa, Oliver Ebenhöh

## Abstract

A thorough understanding of ecosystem functioning in the Arctic Ocean, a region under severe threat by climate change, requires detailed studies on linkages between biodiversity and ecosystem stability. The identification of keystone species with special relevance for ecosystem stability is of great importance, yet difficult to achieve with established community assessments. In the case of microbes, metabarcoding and metagenomics offer fundamental insights into community structure and function, yet remain limited regarding the ecological relevance of individual taxa. To overcome this limitation, we have developed an analytical approach based on three different methods: Co-Occurrence Networks, Convergent Cross Mapping, and Energy Landscape Analysis. These methods enable the identification of seasonal communities in microbial ecosystems, elucidate their interactions, and predict potential stable community configurations under varying environmental conditions. Combining the outcomes of these three methods allowed us to define 38 keystone species in the Arctic Fram Strait that represent different trophic modes within the food web, and might signify indicator for ecosystem functionality under the impact of environmental change. Our research reveals a clear seasonal pattern in phytoplankton composition, with distinct assemblages characterizing the phases of carbon fixation (polar day) and consumption (polar night). Species interactions exhibited strong seasonality, with significant influence of summer communities on winter communities but not vice versa. Spring harbored two distinct groups: consumers (heterotrophs), strongly linked to polar night, and photoautotrophs (mainly Bacillariophyta). These groups are not causally related, suggesting a “winter reset” with selective effects that facilitates a new blooming period, allowing survivors of the dark phase to emerge. Energy Landscape Analysis showed that winter communities are more stable than summer communities. In summary, the ecological landscape of the Fram Strait can be categorized into two distinct phases: a production phase governed by specialized organisms that are highly responsive to environmental variability, and a heterotrophic phase dominated by generalist species with enhanced resilience.

## 1 INTRODUCTION

The Arctic Ocean is a unique ecosystem, undergoing major transitions during climate change. Over the past two decades, temperatures have risen more than twice compared to the global average (Meredith et al., 2019), linked to reduced in sea-ice and snow cover, which exacerbates warming trends. In particular, the extent of Arctic sea ice has declined (Meredith et al., 2019). These environmental changes have a wide range of consequences, including profound shifts in biodiversity (Sala et al., 2000), and thus have a fundamental impact on ecosystems of the Arctic Ocean. There are first signs that the geographical ranges of temperate species are shifting northwards (Kraft et al., 2013), while polar fish and ice-associated species experience a reduction in their habitat due to changing environmental conditions. These ecological changes impact the entire ecosystem stability (Meredith et al., 2019). The complex relationship between biodiversity and ecosystem stability remains poorly understood, particularly in the Arctic Ocean. Consequently, the rapid changes in Arctic sea ice and environmental conditions urgently require an improved understanding of the mechanisms governing the resilience and stability of biological processes and ecosystem functions in the Arctic Ocean. Within marine ecosystems, primary production is a key service supporting all trophic levels (Eppley and Peterson, 1979; Lin et al., 2003), with implications for biodiversity, the abundance and community structure at higher trophic levels, and carbon sequestration. This distinct ecosystem feature is supported by a highly productive microalgal community that thrives in sea ice, accompanied by a remarkably diverse heterotrophic community ranging from bacteria to metazoans (Bluhm et al., 2017). Recent decades have seen a remarkable increase in pelagic phytoplankton and primary production in the Arctic Ocean, a direct consequence of global warming (Arrigo et al., 2015; Lewis et al., 2020; Nöthig et al., 2020).

In the Arctic Ocean (CAO), sea-ice algae rather than phytoplankton account for much of the primary production (Gosselin et al., 1997; Fernández-Méndez et al., 2015)as they have the potential to initiate pelagic blooms beneath the ice (van Leeuwe et al., 2022). Typically, phytoplankton growth starts mainly within the marginal ice zone in spring, co-occurring with increased solar radiation and meltwater-induced stratification (Clement Kinney et al., 2020). Over the past three decades, increasing evidence has documented the occurrence of under-ice blooms in the Arctic Ocean (Strass and Nöthig, 1996; Fortier et al., 2002; Leu et al., 2011; Assmy et al., 2017), while phytoplankton in the water column below the ice shows significant differences from the microalgal communities in the sea ice (Hardge et al., 2017). However, changes in biodiversity key species related to the increase in Arctic pelagic primary production and its impact on the marine ecosystem stability are currently unresolved.

Recent findings indicate that high temperatures in natural ecosystems may affect ecological stability, whereas the consequences of alterations to biodiversity remain variable (Zhao et al., 2023). Nevertheless, the underlying mechanisms remain a subject of debate and limited understanding (Loreau and De Mazancourt, 2013). The presence of nearly 2,000 phytoplankton taxa and 1,000 ice-associated protists in the Arctic (Bluhm et al., 2011) indicates the relevance of identifying keystone species in this wealth of Arctic marine microbial diversity that account for ecosystem stability (Frey, 2017; Barber et al., 2015; Richter-Menge and Farrell, 2013).

Understanding biological and ecological dynamics across seasonal environmental gradients is substantially fostered by novel statistical approaches. In polar ecosystems, these gradients, including Polar day and -night, as well as variations in sea-ice cover, stratification, or nutrient concentrations. Techniques are now accessible to assess the impact of ecological variables on ecosystem stability. For instance, co-occurrence networks (CON) determine and visualize how species coexist within communities or ecosystems (Priest et al., 2023; Ma et al., 2016). However, in natural ecosystems, species interactions are subject to variation as a result of changes in environmental conditions, which can cause a transition from one stable state of co-occurrence to another (Ives and Carpenter, 2007). Cross-convergence mapping (CCM) helps to identify causality of co-occurrence in complex ecosystems, i.e. which organisms might share mutual or other direct relationships. Energy Landscape Analysis (ELA) aids in building ecological models that simulate and predict how ecosystems respond to disturbances or changes of environmental parameters (Sugihara et al., 2012; Suzuki et al., 2020, 2021).

In this study we establish a mathematical methodology to reveal seasonal patterns, suggest causal ecological relationships and identify microbial key species in Western Fram Strait. This major,gateway between Arctic and Atlantic Oceans has been studied for over 20 years within the framework of the Longterm ecological research site HAUSGARTEN and FRAM observatories (Soltwedel et al., 2016). Our study contributes an extended mathematical perspective on microbial inventories in Fram Strait, showing seasonal patterns and the influence of sea-ice on microbial dynamics and the biological carbon pump (Metfies et al., 2017; Wietz et al., 2021; von Appen et al., 2021; Cardozo-Mino et al., 2023; Wietz et al., 2024). Based on a four-year metabarcoding dataset of microeukaryotic taxa in context of rich oceanographic data, sampled year-round in approx. biweekly intervals, we develop scenarios of their long-term resilience. Additionally, we predict taxa that play a crucial role in maintaining stable communities within the Arctic eukaryotic planktonic food web. Furthermore, we seek to define keystone species that can serve as indicators for monitoring the consequences of environmental change for Arctic marine ecosystem stability. Using an unprecedented combination of network analysis techniques like co-occurrence networks and cross convergence mapping, along with energy landscape analysis, our objective is to elucidate which factors might determine the stability of Arctic marine ecosystems. This approach will significantly improve our understanding of the effects of climate change on this ecosystem.

## 2 METHODS

### 2.1 Sampling and Data

Samples were collected with Remote Access Samplers (RAS; McLane) deployed in conjunction with oceanographic sensors over four annual cycles (01.08.2016 to 16.09.2020 (96 Samples)) at the F4 mooring (79.0118N 6.9648E) of LTER HAUSGARTEN and FRAM in the Fram Strait (Soltwedel et al., 2005; Oldenburg et al., 2024). Each RAS contains 48 sterile bags, each collecting water samples of 500 mL at programmed sampling intervals. The samples collected from 2016 to 2018 reflects the pool of up to two samples collected one hour apart in two individual bags. Since 2018, we pooled samples taken 7 to 8 days apart from two consecutive weeks. The samples were preserved by adding 700 μl of mercuric chloride (7.5% w/v) to the bags prior to sampling. Following RAS recovery, water samples were filtered onto Sterivex filter cartridges with a pore size of 0.22 μm (Millipore, USA). Filters were stored at -20°C until DNA extraction and ribosomal metabarcoding of 18S rRNA reads using primers 528iF (GCGGTAATTCCAGCTCCAA) and 926iR (ACTTTCGTTCTTGATYRR).

The resulting amplicon sequence variants (ASVs) were classified using the PR2 4.12 database (see Supplementary - Methods). We normalised raw ASV counts for CON and CCM using the Hellinger transformation but did not for the energy landscape analysis; hence a different normalisation is introduced in the for the rELA implementation (Suzuki et al., 2021).

Temperature, salinity and oxygen concentration were measured with a CTD-O 2 attached to the RAS. Physical oceanography sensors were manufacturer-calibrated and processed as described under (von Appen et al., 2021). Raw and processed mooring data are available at PANGAEA https://doi.org/10.1594/PANGAEA.904565, https://doi.org/10.1594/PANGAEA.940744, https://doi.pangaea.de/10.1594/PANGAEA.941125 and https://doi.org/10.1594/PANGAEA.946447. For chemical sensors, the raw sensor readouts are reported. The fraction of Atlantic and Polar Water were computed following (von Appen et al., 2018) for each sampling event and reported along with distance below the surface (due to mooring blowdown). Sea ice concentration derived from the Advanced Microwave Scanning Radiometer sensor AMSR-2 (Spreen et al., 2008) were downloaded from the Institute of Environmental Physics, University of Bremen (https://seaice.uni-bremen.de/sea-ice-concentration-amsr-eamsr2). Sentinel 3A OLCI chlorophyll surface concentrations were downloaded from https://earth.esa.int/web/sentinel/sentinel-data-access. For all satellite-derived data, we considered grid points within a radius of 15km around the moorings. Surface water Photosynthetically Active Radiation (PAR) data, with a 4 km grid resolution, was obtained from AQUA-MODIS (Level-3 mapped; SeaWiFS, NASA) and extracted in QGIS v3.14.16 (http://www.qgis.org).

We considered eight environmental variables : mixed layer depth (MLD in m), water temperature (temp °C), polar-water fraction (PW frac %), chlorophyll concentration from in situ sensor (chl sens *μ*g *l*^−1^), PAR (*μ* mol photons *m*^−^2*d*^−^1), Salinity (PSU), oxygen concentration (O2 conc *μ*mol *l*^−1^) and sampling depth (depth m) (von Appen et al., 2021),.

### 2.2 Co-Occurrence Network

The abundance of species over the full observation period were converted into temporal profiles by employing Fourier transformation techniques to time-series signals. These temporal profiles rely on the 14 Fourier coefficients. We chose 14 coefficients because they reflect the majority of observed species abundance peaks within the four years. To investigate the similarity of temporal profiles between species pairs, we performed pairwise correlations between the individual temporal profiles, where pairs with higher Pearson correlation value show also a similar temporal profile. Pairs with at least 0.7 (p*<*0.05) Pearson correlation were then visualized in an undirected graph. Only positive correlations were retained to later focus on co-operative relationships. To identify strongly connected components that reflect the existing communities of co-occurring species, we applied the Louvain community detection algorithm (Blondel et al., 2008) on the entire graph. The entire process was implemented using the CCM and networkx packages in Python; visualization was performed using Cytoscape with the Edge-weighted Spring-Embedded Layout (Shannon et al., 2003). The whole co-occurrence network construction is described in Supplementary - Methods Co-Occurrence Network.

#### 2.2.1 Distance between Clusters

To measure the distance between previously defined Louvain communities (clusters), we applied UMAP on time-series signals obtained after Fourier decomposition of the abundance data. From this we generated a three-dimensional embedding space. Centroids for each cluster were calculated within this space (see Supplementary - Results Figures S4, S5, S6). The network distance between clusters was determined as the Euclidean distance between their centroids. Subsequently, a distance matrix was created and distances were rounded to integers, with only significant connections retained.

### 2.3 Convergent Cross Mapping

Convergent Cross Mapping (CCM) identifies potential causal relationships between variables in time series data. It quantifies how knowledge of the time series of one species allows predicting the time series of an another species. We first built a CCM network from all pairwise combinations. From this, we extracted the in- and outgoing edges between nodes that are also connected in the co-occurrence network. We used the implementation of Normalized Mutual Information (NMI) from https://github.com/polsys/ennemi by Petri Laarne and the Convergent Cross Mapping by Implementation from Javier, Prince https://github.com/PrinceJavier/causal_ccm (Javier et al., 2022) to measure the strength of the causal relationship considering also non-linear relations. We could show that the implementation of Normalized Mutual Information (NMI) results into similar findings as the original implementation based on Pearson correlation (Veilleux, 1979; Sugihara et al., 2012) (see Supplementary - Methods).

Using a permutation approach (Ma et al., 2016) on the connectivity of the network we calculated significance values for the edge weights, quantifying whether the respective NMI values are greater than expected for random edges (see Supplementary - Results). The whole CCM network construction and validation are described in Supplementary - Methods and Supplementary - Results (Convergent Cross Mapping) of the Supplementary Information.

#### 2.3.1 Aggregation on cluster level

We simplify the network of interactions between single species into a network of interactions between clusters. For this, we assign a weight to a directed edge between two clusters by calculating the arithmetic mean of NMI of all (directed) edges connecting species belonging to the respective clusters. This process effectively reduces the number of items in the node cloud, representing clusters through a unified composite node.

### 2.4 Energy Landscape Analysis

Energy Landscape Analysis is a method based on statistical physics. From data for many points in time, which contain species abundance and environmental variables, an *energy landscape* is reconstructed. This energy landscape is a function that maps ASV abundance and environmental variables to an energy value. In analogy to the potential energy in physics, a (local) minimum of this energy landscape indicates a stable community state. Here, we reconstruct the energy landscape function based on the complete time series of ASV abundance together with the available environmental data. We use the reconstructed function to determine the stability of observed communities, and in particular the seasonal clusters determined by the co-occurrence network, and we predict the most stable community compositions. Moreover, because this function also depends on environmental variables, we can predict how stable a given community is under perturbed environmental conditions. Assessing stable community states and how they change across environmental shifts is crucial for comprehending the resilience and adaptability of ecosystems in the face of environmental challenges. Our analysis focused on the Top 100 most abundant ASVs within each cluster. Outliers were excluded solely for the purpose of plotting. Details of our analysis, including parameters and thresholds applied, are described in in Supplementary - Methods (Energy Landscape Analysis). Understanding the existence and the nature of stable community states and how they change in response to environmental shifts is crucial for comprehending the resilience and adaptability of ecosystems in the face of various ecological challenges. Details of our analysis, including parameters and thresholds applied, are described in in Supplementary - Methods (Energy Landscape Analysis). Our analysis focused on the Top 100 ASVs within each cluster. Outliers were excluded solely for the purpose of plotting.

### 2.5 Keystone Species Definition

After collecting attributes from a co-occurrence analysis and distinguishing between potential ecological influence and occurrence just by chance, we calculated the stable states for different clusters using ELA. This information was merged to suggest potential keystone species. We defined a keystone species as an ASV with i) a significant influence on other organism in the network (significant NMI value), ii) a high centrality (closeness) value within its co-occurrence community and iii) presence in at least one stable state as predicted by ELA. A significant high centrality value was determined by comparing each centrality value of a single node to the average centrality values of all nodes from the graph using a one sided, one-sample t-test with Benjamini-Hochberg correction for multiple testing (similar to (Guimera and Nunes Amaral, 2005; Joyce et al., 2010).

### 2.6 Season Definition

For assessing results in context of the entire annual variability over which samples were collected, we defined the seasons as follows, based on month and the availability of light (PAR). In the case of Cluster 01TA, the maximum month is August. However, several nodes are also present in September and October. Consequently, we mapped this cluster to the autumn season, in order to model a transition from the autumn cluster.

## 3 RESULTS

We examined a dataset of 1,019 eukaryotic ASVs and eight environmental parameters compiled over four years at mooring site F4 in the West Spitsbergen Current (WSC) in Fram Strait. The aim was to characterize species communities, analysing causal relationships between ASVs and to identify keystone and resilient species with respect to the impact of various environmental conditions.

To accomplish this, we established a novel computational pipeline, coupling co-occurrence analysis with convergence cross mapping and energy landscape analysis. This allowed us to identify causal interactions among species in a co-occurring community and to identify stable community states across different environmental conditions.

### 3.1 Co-Occurrence Network reveals seasonal dynamics

The co-occurrence network (CON) comprised eight connected components, with a major component accounting for 98% (935) of all nodes, which are connected by 8,610 edges. In the following, we focus on this major connected component. The resulting undirected graph notably displays a clear seasonal cyclic pattern (Figure 2 A).

**Figure 1.**
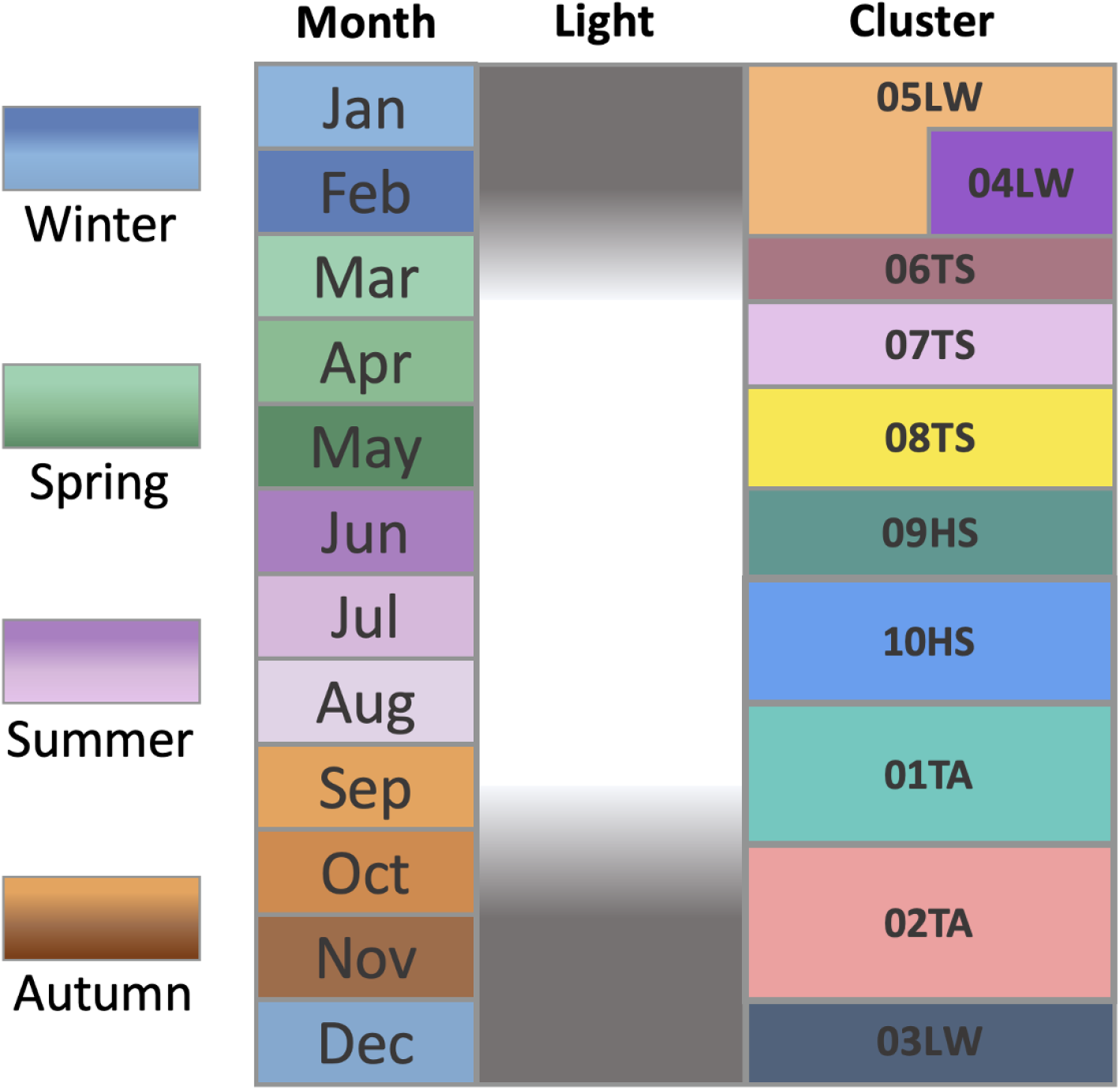
(Schematic) Cluster definition: Names are based on light availability (as defined in Oldenburg et al. (2024)), categorized in transition areas between dark and light (T), high light (H) and low light (L) phases based on PAR parameter (Figure 4) and the season spring (S), summer(S), autumn (A) and winter (W).

**Figure 2.**
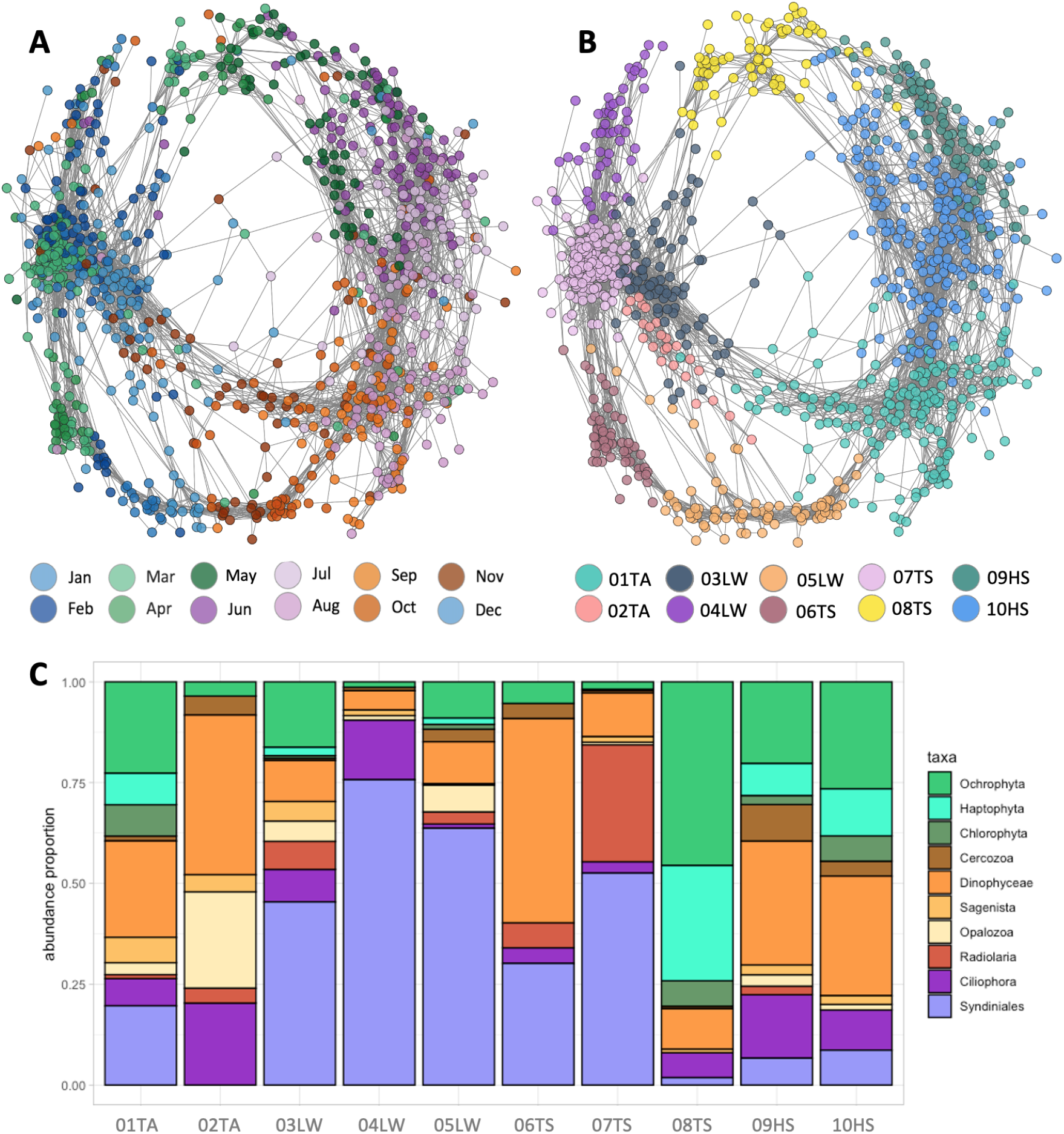
Co-Occurrence Networks and microeukaryotic composition at mooring F4 from 2016-08-01 to 2020-09-17. Each network node represents an ASV, and each edge represents a similar temporal pattern of two ASVs. The edge weights correspond to the Pearson correlation coefficients determined from the comparison of the individual ASV temporal profiles. ASVs are connected if the coefficient is *r >* 0.7, *p <* 0.05. **A:** Node color reflect the month in which the ASV exhibit maximal abundance, calculated from the maximum abundance mode for each year ranging from January to December. **B:** In this representation, nodes are coloured based on the community membership that was determined by the Louvain community detection algorithm. **C:** The relative abundance of the top 10 taxonomic classes by cluster (‘HS’ high light summer, ‘LW’ low light winter, ‘TS’ corresponds to transition spring and ‘TA’ transition autumn). Colour shades illustrate the assignment to auto-(green), mixo-(orange) or heterotroph (purple) .

The network was partitioned using the Louvain community detection algorithm (Blondel et al., 2008), revealing ten discrete community clusters (Figure 2 B) labeled by the season in which the majority of clusters members had their maximum abundance (Table 1). To further group the clusters, we submerge each three month period to one season. Two clusters were assigned to the transition autumn period (01TA and 02TA), three clusters were associated with the low light winter period (03LW, 04LW and 05LW) and three clusters with the transition spring period (06TS, 07TS and 08TS). Finally, clusters 09 and 10 were allocated to the high light summer season (09HS, 10HS).

**Table 1.** Co-occurrence network clusters. Ten labeled clusters with their assigned season based on month in which the ASV exhibit maximal abundance, calculated from the maximum abundance mode (majority vote) for each year ranging from January to December, the number of ASVs per cluster, and the number of significant edges in network graph. The stats denote that the MaxMonth is in August, but several nodes are also in September and October, therefore we mapped it to autumn, to model a transition autumn cluster.

| Cluster | MaxMonth | Season | Number of ASVs | Number of edges |
| --- | --- | --- | --- | --- |
| 01TA | Aug* | Autumn* | 162 | 997 |
| 02TA | Nov | Autumn | 24 | 86 |
| 03LW | Dec | Winter | 78 | 522 |
| 04LW | Feb | Winter | 52 | 212 |
| 05LW | Dec | Winter | 74 | 467 |
| 06TS | Mar | Spring | 51 | 388 |
| 07TS | Mar | Spring | 153 | 3040 |
| 08TS | Apr | Spring | 63 | 262 |
| 09HS | Jun | Summer | 88 | 509 |
| 10HS | Jul | Summer | 190 | 1189 |

### 3.2 Community composition

We explored the taxonomic community per cluster to explore the seasonal associations of each specific taxonomic group. Alpha biodiversity, measured by Shannon entropy, decreases from summer through autumn and winter, gradually decreasing towards spring (Supplementary - Results Figure S1). The beta biodiversity, measured by Bray-Curtis distance (Supplementary - Results Figure S2), between the winter and spring clusters (03LW, 04LW, 05LW and 06TS) is notably lower than the most other scores, except that between 01TA and 10HS. Cluster 02TA exhibits on average a higher beta diversity compared to all other clusters, which can be explained by the fact that 02TA is the smallest cluster in terms of the number of ASVs (Supplementary - Results Figure S2).

In our study, we found distinct taxonomic compositions within various clusters. Photosynthetic organisms like Ochrophyta and Haptophyta dominate the light phases (Oldenburg et al., 2024). In late spring (cluster 08TS) phototrophs make up more than 75% of ASVs, while during summer (clusters 08TS and 09HS) and early autumn (10HS) they still comprise over 25% of all ASVs. Mixotrophs are highly abundant in most clusters, while they clearly dominate during the late autumn transition (cluster 02TA). Through the complete dark period (clusters 03LW, 04LW and 05LW) as well as in early spring (06TS and 07TS), heterotrophs, particularly Syndiniales, are dominant wiht a clear peak of abundance (more than 90% of ASVs) in mid winter (cluster 04LW). During early spring (cluster 06TS) when sunlight appears again, mixotrophs increase in their abundance, highlighting the nuanced trophic dynamics during the annual cycle. This comprehensive analysis at taxa level provides insights into the composition of these clusters, shedding light on the prevalence and distribution of specific classes within distinct seasonal communities (Figure 2 C).

### 3.3 Convergent Cross Mapping identifies Community Interactions

Convergent Cross Mapping (CCM) was applied to predict causal relationships within and between seasonal clusters based on the underlying ASV dynamics. We project the CCM-derived weights onto the co-occurrence network, resulting in a directed graph consisting of 17,220 directed edges and 935 nodes. Here, a directed edge indicates that knowledge of the dynamics of the source node allows predicting the dynamics of the target node.

A comparative analysis of edge weights within the CCM network was conducted. The connectivity derived from the co-occurrence network was compared with theoretical edge weights and randomly permuted connections. To perform this comparison, a two-sided Kolmogorov-Smirnov test was used. The theoretical edge weights were derived from all possible connections between pairs of nodes in the co-occurrence network, excluding the existing true links (Section 3.3). The findings clearly show that there is a stronger causal influence (higher NMI values) between co-occurring species compared to random or to unconnected nodes (Supplementary - Results Figure S8 and S9 and Table S1 and S2). The significance level was set at a nonparametric p-value of less than 0.05 calculated similar to Ma et al. (2016). Trimming edges with non-significant NMI values (Supplementary - Results Figure S10), produced a network graph consisting of 4,597 edges and 719 nodes, divided into 18 disconnected components, with the largest module encompassing 706 nodes with 4,572 edges(Fig. 3 A,B).

**Figure 3.**
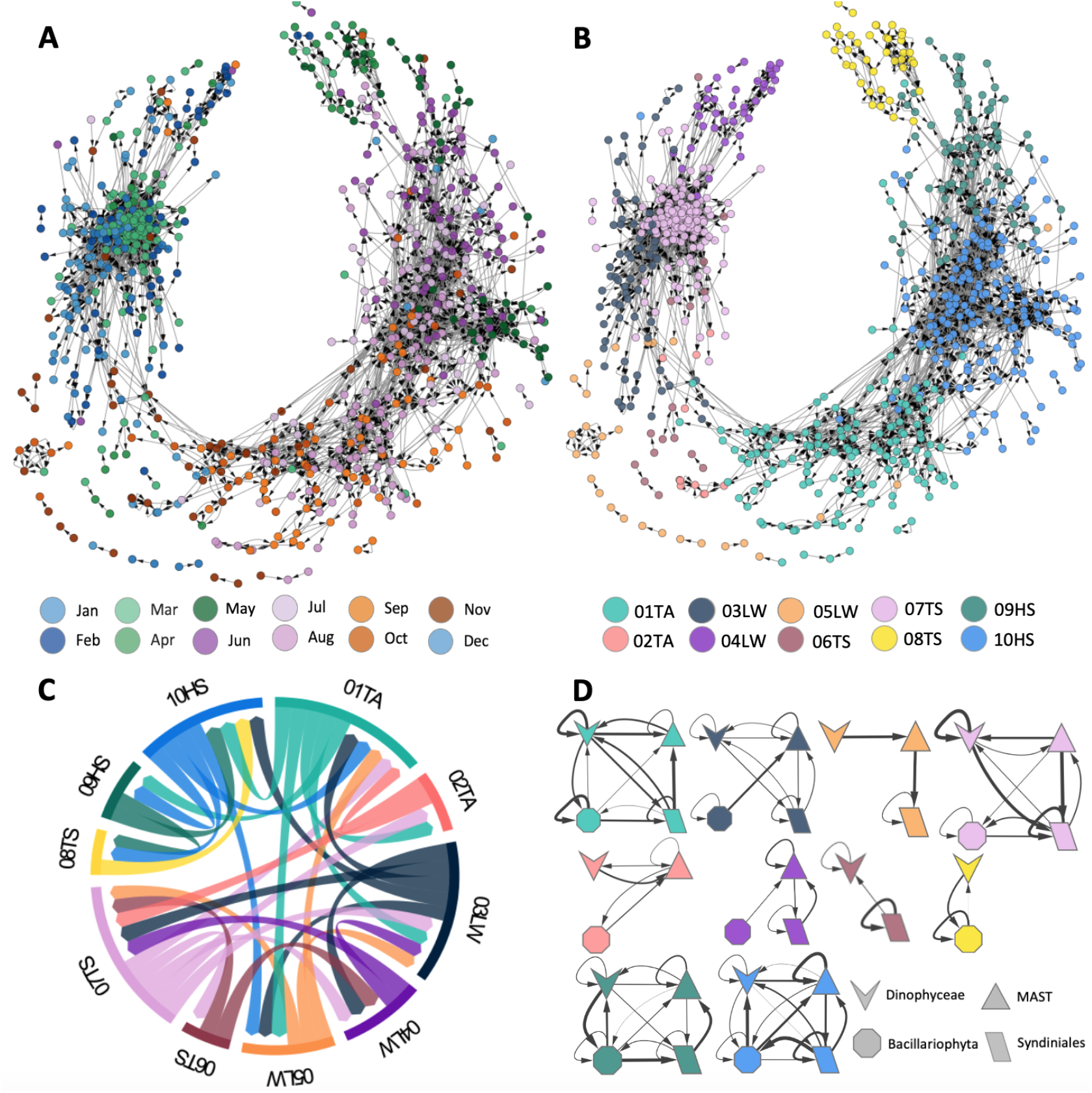
Convergence Cross Mapping Networks of microeukaryotes at mooring F4 from 2016-08-01 to 2020-09-17. Each node in the CCM network represents an ASV, and each edge represents the causal influences. The edge weight corresponds to the Normalized Mutual Information determined from the comparison of the individual ASV and their predicted representation in the shadow manifold. ASVs are connected if the smoothed p-value of the weight is *p <* 0.05. **A:** Node color reflects the month in which the ASV exhibits maximal abundance, calculated from the maximum abundance mode for each year ranging from January to December. **B:** In this representation, nodes are coloured based on the community membership that was determined by the Louvain community detection algorithm. ‘HS’ labels denote high light summer, ‘LW’ represents low light winter, ‘TS’ corresponds to transition spring, and ‘TA’ indicates transition autumn. **C:** The Normalised Mutual Information aggregated across the edges between the clusters, visually represented by thickness of the arrows corresponding to their respective values. Corresponding colour visually represent the clusters. **D:** Interaction analysis between taxonomic clusters. For each of the ten clusters, the interactions between ASV groups are examined at class level, considering ‘Syndiniales’, ‘Dinophyceae’, ‘Bacillariophyta’ and ‘MAST’. The cluster assignments are marked by different colors. The thickness of the arrows denotes the strength of the interaction, while the shapes represent the various taxa groups at the class level.

Of the total 12,648 edges eliminated during trimming, 18.16% represent connections between species reaching their peak abundance in March. This selective removal has a profound impact on the network structure, (Figure 3 A). In contrast to the co-occurrence network, the causal interaction network “breaks” during spring season, as demonstrated in Figure 3 A (months March and April). The winter cluster 05LW and the spring cluster 06TS collapse (see Figure 3 B), meaning that nodes disintegrate and large parts of the clusters are no longer connected to the rest of the network. This suggests that the corresponding connections in the co-occurrence network are not a result of causal interactions, but rather result from other factors, possibly caused by the prevailing environmental conditions. Analysis of the betweenness centrality reveals that species of Picozoa, Leegaardiella, Acantharea, Dinophyceae, MAST-1, and Syndiniales serve as essential hub nodes throughout the seasonal cycle in the network, highlighting their crucial function in maintaining network stability.

### 3.4 Community Interaction

Analyzing cluster interactions revealed distinct patterns. We measure distance of clusters by “network distance”, a metric designed to evaluate the separation between clusters. This measure is computed by assessing the distance between the centroids of clusters within the dimension-reduced UMAP embedding space (Supplementary - Results). Distances between clusters thus determined range between one and seven. Proximate clusters (a network distance of two to four), exhibited notably higher connectivity compared to clusters situated further apart (distance five to eight). Figure 3 C provides a visual and quantitative representation of these interactions. Subsequent analysis revealed a prevalence of connections at a network distance of two (91.5% of total connections), followed by distances of three (7.4% of total connections), and four (0.7% of total connections) (Supplementary Results Figure S4, S5, S6).

For each seasonal cluster, we investigate in detail the mutual influence (NMI) of four taxonomic groups selected from the top ten classifications (Figure 2 D): Bacillariophyta, Syndiniales, Dinophyceae, and MAST (all MAST-X variants were classified under MAST). Bacillariophyta primarily comprises photo-autotrophic species (Mann et al., 2017), while Syndiniales include parasitic species, most of them characterized by their heterotrophic lifestyle (Suter et al., 2022). Dinophyceae are known for their diverse array of species and ecological roles, from symbionts to planktonic autotrophs (Lin et al., 2022). MASTs are heterotrophic protists and contribute substantially to protist abundances in the ocean. They play a crucial role in marine ecosystems being among the dominant eukaryotes in the Arctic Ocean (Thaler and Lovejoy, 2014; Lin et al., 2022).

These four taxonomic groups are primarily distinguished by their unique lifestyles and ecological roles as primary producers, consumers, parasites, or endosymbiotic interactors. These distinctions form the basis for our analysis of their contributions to the ecosystem. Hence, by summarizing the members of each group into single nodes, we analyzed their cross-interactions using the information obtained from the CCM network.

The strength and direction of interactions between these taxonomic groups varied over the annual cycle (see Figure 3 D). During the spring-summer and summer-fall transition, clusters 02TA and 06TS displayed fewer and weaker connections compared to other clusters (see Figure 3 B). This suggests dynamic changes in community structure during these transition phases, with ecological interactions between individual species either yet to be established or no longer present. The start of the polar night (cluster 05LW), we detected the most substantial influence from the dinoflagellates (shown by the thickest arrow in Figure 3 D) to the pico-eukaryotic heterotrophic groups Syndinales and MAST, suggesting a crucial ecological role of dinoflagellates for the establishment of the winter community. Overall, the strength of the links between the taxonomic groups decreases as the polar night progresses and has its minimum at the peak of the polar night in December (Cluster 04LW and 03LW). The lack of strong connections between taxonomic groups during the deepest polar night indicates that there are only a few ecologically significant interactions within the microeukaryotic community.

During polar day, the connections between taxonomic groups become stronger and reach their peak during the zenith of the polar day (Cluster 10HS). Notably, Bacillariophyta (i.e. diatoms) showed the most robust connections during the polar day, owing to their role as predominat phototrophic biomass producer and form the foundation of the marine food web. However, towards the end of the growth period, the impact of MAST on Dinoflagellates becomes more pronounced (as shown by the thicker arrow), indicating an essential involvement of this pico-eukaryotic heterotroph in the ecosystem during the late polar day; a signature of the transition from primary production to recycling.

### 3.5 Community and Environment Interactions

For a more detailed understanding of which environmental conditions align with seasonal community clusters, we conducted a correlation analysis (Figure 4). Cluster 10HS displayed a significant positive correlation with Photosynthetically Active Radiation (PAR) (0.64) and temperature (0.52), but a significant negative correlation with Mixed Layer Depth (MLD) (-0.64). This cluster thrives in environments with high light and temperature levels but less deeper mixed layers. Cluster 03LW exhibits the opposite behavior, showing a moderately positive correlation with MLD (0.25) and polar water fraction (PW frac) (0.21) while displaying an inverse relationship with PAR and temperature (-0.35 and -0.37, respectively).

**Figure 4.**
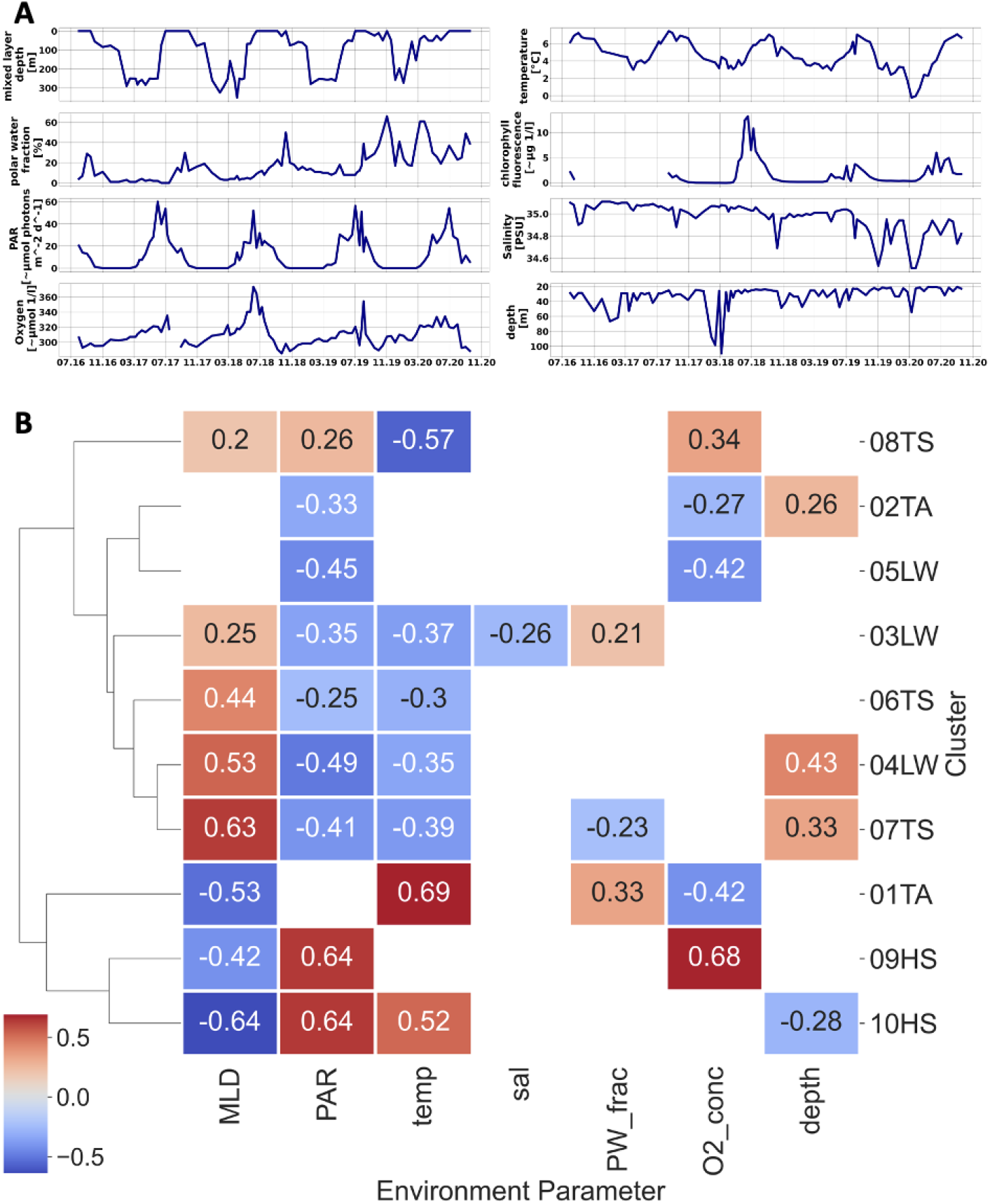
Environmental data and their correlation with Louvain clusters. **A):** Environmental data for F4 from 2016-08-01 to 2020-09-17. The x-axis represents the time period, while the y-axis indicates the following parameters: Mixed Layer Depth (MLD) [m], Temperature [°C], Chlorophyll Fluorescence [*μ* g/L], Polar Water Fraction [%], Photosynthetically Active Radiation (PAR) [*μ* mol photons/*m*^2^/d], Salinity [Practical Salinity Units (PSU)], Oxygen Concentration [*μ* mol/L], Depth of measurement [m]. **B): Correlations between environmental parameters and seasonal Louvain clusters**. The displayed chart shows the environmental parameters from section a) in relation to seasonal clusters. These clusters are characterized by the cumulative relative abundance of ASVs. ‘TA’ denotes transition autumn, ‘LW’ represents low light winter, ‘TS’ transition spring and ‘HS’ high light summer. The colour gradient used in the heatmap illustrates the strength of the correlation visually, with blue shades indicating negative correlations and red shades the positive correlations. It is worth noting that a significance mask has been applied to show only correlations that are statistically significant.

### 3.6 Energy Landscape Analysis determines stability of microbial communities

Energy landscape analysis assessed the stability of communities under the prevailing environmental conditions. We focus on four clusters representing the four seasons (01TA for autumn, 03LW for winter, 08TS for spring and 10HS for summer). For each of these clusters, we determined the energy landscape, which is a highly complex function that depends on the abundances of all ASVs and the environmental parameters (see Figure 5). To approximately visualize this landscape, we plot an interpolated smooth surface as a function of the two most significant NMDS dimensions. In addition, for each time point, we evaluate the energy landscape function and represent each energy value by a point in the three-dimensional diagram, where the *z*-axis represents the energy value.

**Figure 5.**
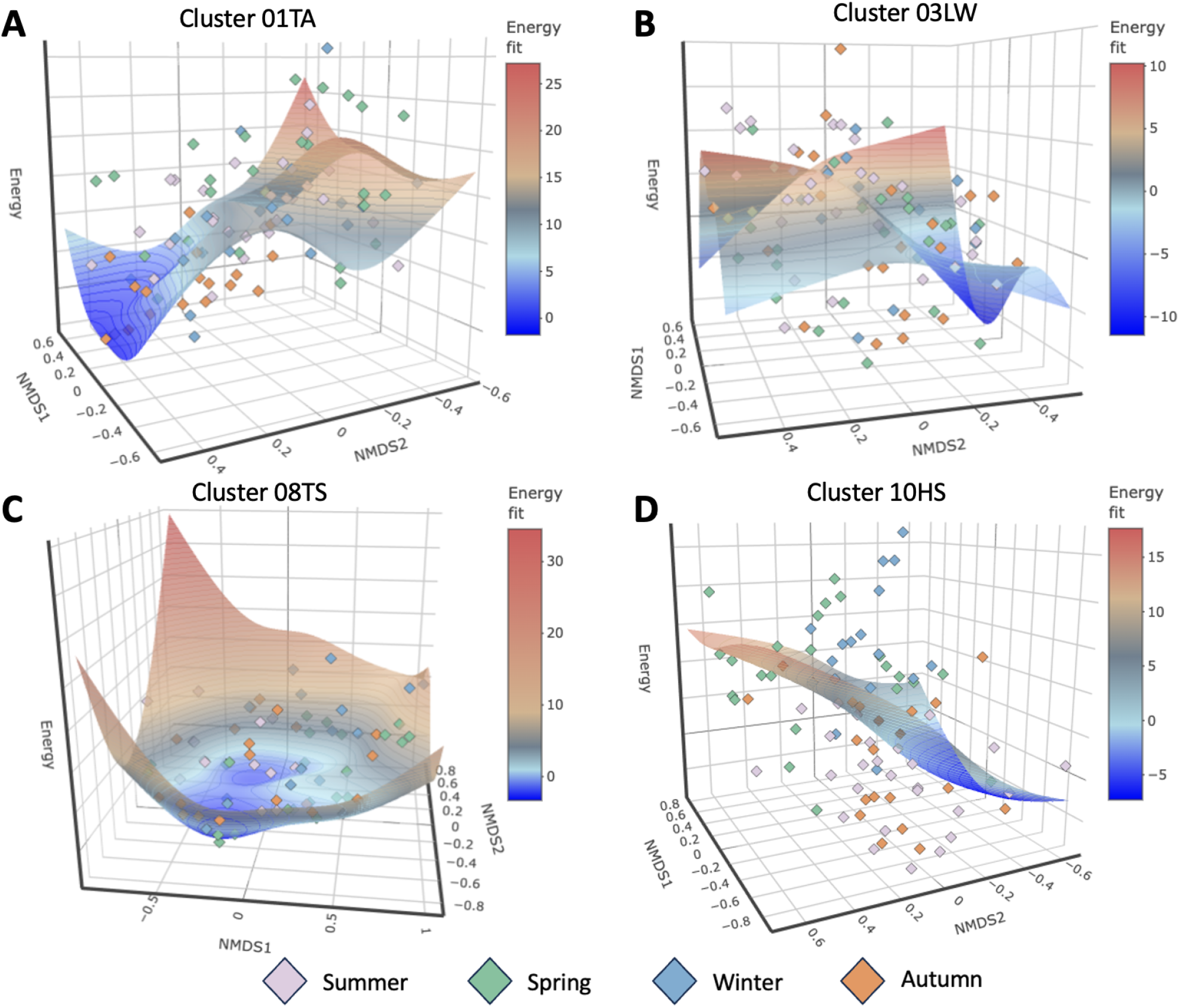
Energy landscapes depicting community structure dynamics. The plots display the reconstructed energy landscape on the NMDS surface for a cluster of each season. Environmental landscapes over the NMDS surface are reconstructed for each of the four example clusters. The z-axis displays the energy, while the x- and y-axes display the first and second NMDS dimensions. The landscape contours were estimated using a smoothing spline approach with optimized penalty parameters. Community states, which are defined by ASV compositions and occupy lower-energy regions, indicate higher stability within the energy landscapes. **A):** The transition autumn cluster 01TA. **B):** The low light winter cluster 03LW. **C):** The transition spring cluster 08TS. **D):** The high light summer cluster 10HS.

For the landscape reconstructed for cluster 01TA (Figure 5 A) the community displayed lower energy values than in other seasons; demonstrating high stability of the autumn community. This demonstrates that the autumn communities exhibit a high stability. For the winter cluster 03LW (Figure 5 B), the picture is less clear. Whereas the interpolated energy landscape has a more pronounced minimum, the energy values of the observed communities are not clearly separated. As a tendency, the summer communities have a high energy value, demonstrating that summer communities are unstable in winter conditions. However, spring and autumn communities exhibit comparable energy values as winter communities, which indicates that stable community structures in winter conditions are not clearly defined. This trend is even more pronounced for the spring cluster 08TS (Figure 5 C). Here, the interpolated energy landscape shows a broad and shallow minimum, and the energy values of the all observed communities, regardless of the season in which they are found, are very similar. This suggests that under spring conditions community structures are not very stable and that community compositions show a high plasticity. As a consequence, many different communities may exist under spring conditions. The findings demonstrated that knowledge of the composition of winter communities does not allow for the prediction of the composition of spring communities, in conjunction with the observation of the CCM analysis and the gap between winter and spring clusters (Figure 3 B). Finally, the energy landscape in summer (Figure 5 D) shows a pronounced minimum, in which the observed summer communities are also found. This indicates that summer conditions support well-defined communities with a high degree of stability.

### 3.7 Predicting keystone microeukaryotes in the Fram Strait

According to our definition (Section 2.5) a keystone species is highly connected in the co-occurrence network, has a high influence on other species, and appears in a stable community. By contextualizing the evidence from CON, CCM and ELA, we predict 38 keystone species across the annual cycle within the measured environmental profile (Table 2). 14 of these keystone species are associated with summer clusters (three and eleven are found in clusters 09HS and 10HS, respectively), 13 with winter (eleven and two in the winter clusters 03LW and 04LW, respectively), eight are associated with autumn (cluster 01TA) and three with spring (cluster 07TS). The nine keystone species from the summer clusters belong to the taxonomic groups Ochrophyta (6), Dinophyceae (4),Ciliophora (3) and Cryptophyta (1). These groups include Fragilariopsis, Pseudo-nitzschia, and Thalassiosira, major diatom taxa during Arctic blooms (Von Quillfeldt, 2000) that also serve as prey for microzooplankton (Cleary et al., 2016; Yang et al., 2015). Notably, Fragilariopsis and Thalassiosira exhibited the highest abundance within this cluster. The keystone species in the winter clusters comprise Syndiniales (10), Radiolaria (1), Ochrophyta (1) and Dinophyceae (1), autumn cluster keystone species are Syndiniales (3), Ochrophyta (2), Chlorophyta (1), Dinophyceae (1) and Eukaryota uc (1). The spring keystone species belongs to Syndiniales (1) and Radiolaria (2); reflecting the major ecological strategies including primary production, heterotrophy, and parasitism. The finding of only a few spring keystone species aligns with the greatest variability as shown by ELA (Section 3.6). The emergence of Chlorophyta during early autumn suggests a shift in primary production from Ochrophyta to Chlorophyta, including taxa that may prefer colder temperatures ((Tragin and Vaulot, 2018) and are better adapted to nutrient limitation (Maat et al., 2014).

**Table 2.**
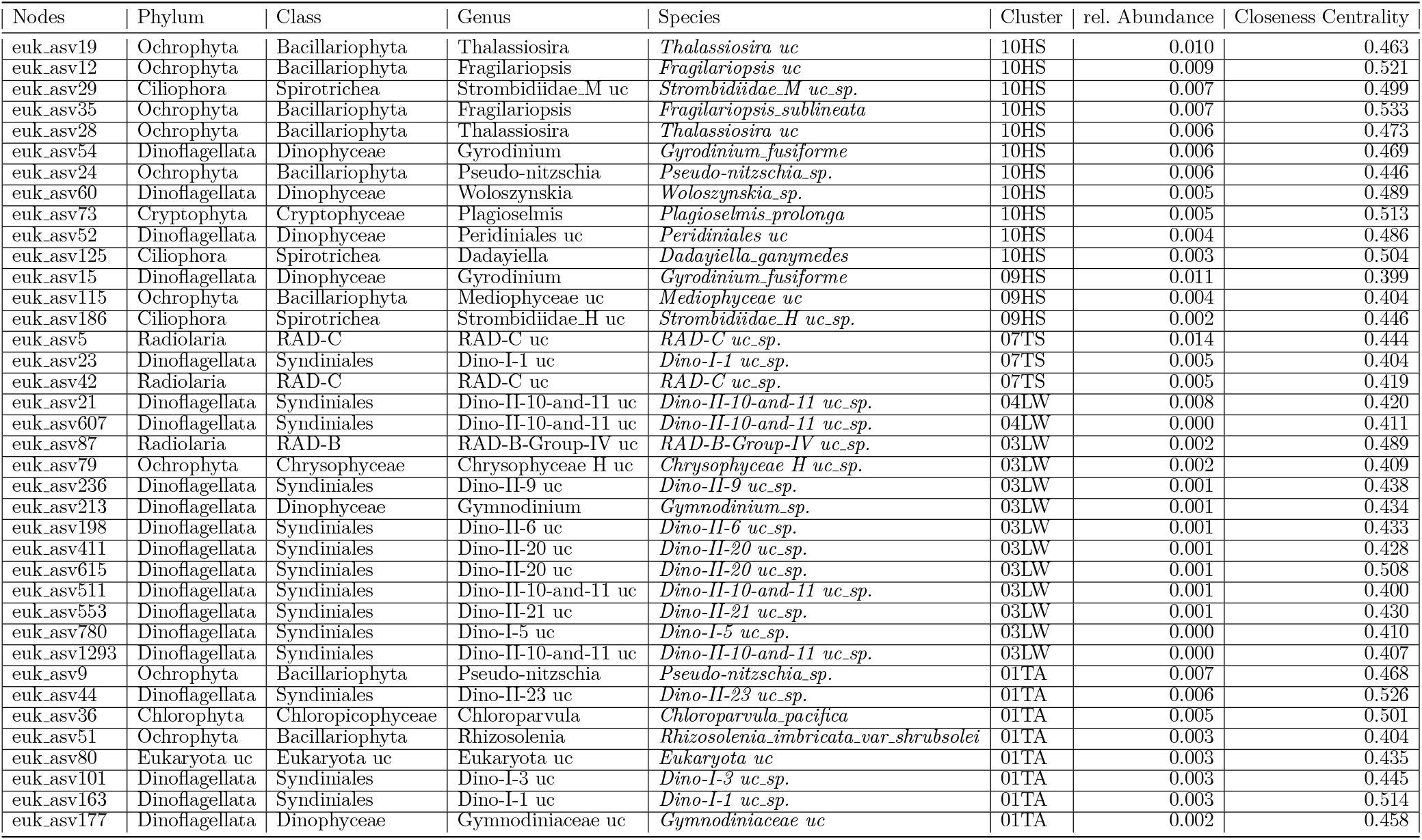
ASV identified as potential keystone species for clusters 10HS, 06TS, 03LW and 01TA. The taxonomic classifications, clusters, raw abundance, proportion of total raw abundance, cluster abundance, and proportion of cluster raw abundance are presented in summary form over the 4-year observation period. The column, significance, indicates if this ASV (Nodes) has at least on significant CCM connection measured in Normalized Mutual Information.

### 3.8 Predicting community stability in altered environments

One critical ecological question is how marine microbial communities change if they experience changing environmental conditions, such as those resulting from climate change. For a first approximation to the potential fate of herein defined clusters under changing conditions, we employed ELA to assess the stability of observed communities under environmental conditions different to those they usually experience. This is done by evaluating the ELA function (see Section 2.4) for given communities and environmental parameters. First, we test the reliability of this approach by evaluating the communities of the summer (10HS) and winter (03LW) clusters under “typical Atlantic” summer or winter days, defined by the averages environmental parameters during a three-month period (see Supplementary - Methods Table S0). As expected the energy values remain almost unchanged when evaluating the summer cluster communities in a typical summer day and the winter cluster communities in a typical winter day (see Supplementary - Results Figure S13). This is to be expected, because the ASVs in the summer clusters assume their most stable community configuration during the summer months, and likewise for the winter cluster. When evaluating the cluster communities under the opposite conditions, i.e. the summer cluster communities with environmental parameters representing a winter day and vice versa, the energy values are drastically changed (see Supplementary - Results Figure S13). Those communities of the summer cluster, which were still present in the winter, exhibit an increased stability (lower energy) than those present in summer. The Atlantic projection panel (Supplementary - Results Figure S13 C and E) display almost opposite oscillatory time courses. Placing communities of the winter cluster in summer conditions has a more differentiated effect. The communities found in winter 2018 and 2020 are highly unstable in this environment, while communities found in summers appear to exhibit an increased stability. This observation allows speculating that some ASVs of the winter cluster might even benefit from warmer summer conditions.

We next investigate a scenario that might result from future temperature increase and accelerated sea-ice melt. Ice-free summers in the Arctic will allow Atlantic water, and with this Atlantic microbial communities, to enter polar regions. We therefore explored the stability of communities from the summer (10HS) and winter (03LW) clusters changes if they experience typical Arctic conditions. For this, we defined “typical” Arctic summer and winter days by selecting extreme environmental parameter values from the central Arctic (see Supplementary - Methods Table S0). Subsequently, we evaluate the energy landscape functions as above with the respective environmental parameters. Interestingly, simulating communities of the summer cluster under Arctic summer or winter conditions results in similar energy values as placing these communities in Atlantic winter days. Specially, communities found in summer are highly unstable whereas communities present in winter show, as a tendency, increased stability (Supplementary - Results Figure S13 E,G,I - left panel, bottom three), indicating that summer cluster communities will face challenges to adapt to Arctic conditions. When comparing the ASVs, which are found in at least one community configuration predicted as stable, we observe that ASVs stable in Arctic environments are almost completely different from those stable in their natural (Altlantic) environment (Supplementary Results Figure S14, Table S9 and S10). To estimate the importance of the ASVs identified to be stable under natural and Arctic conditions, respectively, we determine their average closeness centrality (Supplementary - Results Figure S15). Clearly, ASVs stable in their natural conditions (and in Atlantic conditions) display a significantly (p-value = 2x10^−2^) higher centrality than average, while those predicted to be stable under Arctic conditions show a reduced centrality. Likewise, the average NMI score for outgoing edges, indicating the mean influence of an ASV on other ecosystem members, is higher than average for ASVs stable under natural (and Atlantic) conditions (p-value = 4x10^−2^), but lower than average for ASVs predicted to be stable under Arctic conditions. The reduced connectivity and weaker influence on other ecosystem members suggests that species of the summer cluster, when exposed to Arctic conditions, will play a less important role in determining ecosystem dynamics and stability.

Simulating the effect of Arctic conditions on winter cluster communities reveals that neither communities present in summer nor winter display a pronounced stability. The only exception appear to be communities during spring 2020. Remarkably, this period was characterized by unusually low temperatures (Figure 4 A). In general the ASVs identified as members of stable communities in the original conditions, Atlantic winter, or Arctic winter show a four times higher overlap compared with their summer conditions. These observations suggests that communities of the winter cluster might easily adapt to colder Arctic conditions, which is also supported by the observation that closeness centrality and averaged NMI scores (for outgoing edges) are almost unchanged for these three groups of organisms.

## 4 DISCUSSION

In this study, we developed a new method for investigating ecological time series data based on 18S metabarcoding derived abundance information, by combining three data analysis methods: Co-Occurrence Networks (CON) (Stephens et al., 2009), Convergent Cross Mapping (CCM) (Sugihara et al., 2012), and Energy Landscape Analysis (ELA) (Suzuki et al., 2021). Integrating these three methodological approaches, we aimed to to predict and characterize abundance of keystone microeukaryotes in the West Spitsbergen Current across different seasons, environmental parameters and in relation to other organisms. In addition, we examine how these keystone species are affected by changing environmental conditions, providing insights into potential responses to Arctic warming and Atlantification. We also investigate how different taxa groups affect other taxa groups, and how their effects vary with seasonal shifts and environmental factors.

Our co-occurrence network based on Fourier decomposition differs from previous methods that rely directly on the raw time series signals (Ma et al., 2016; Lima-Mendez et al., 2015; Ma et al., 2020). Our CON based on Fourier decomposition differs from previous methods that rely directly on the raw time series signals (Ma et al., 2016; Lima-Mendez et al., 2015; Ma et al., 2020). The resulting network accurately captured seasonal states and transitions, revealing community clusters that reflect the prevailing community structure (Dunne et al., 2002)): in spring (cluster 08TS) primary producers such as Bacillariophyta appear, and remain throughout the summer (09HS, 10HS), while mixotrophs increase in autumn (01TA, 02TA, 03LW) until an almost exclusively heterotrophic and parasitic taxa dominate in winter (04LW, 05LW, 07TS). The considerable difference of spring clusters to other seasonal clusters (Supplementary - Results Figure S2). can be explained by the rapid environmental changes during this period (i.e. change from darkness to constant daylight within 20 days). The predominance of dinoflagellates in the intermediate phases of spring and autumn indicates that these mixotrophic organisms play a crucial role during transition phases (Jassey et al., 2015; Bruhn et al., 2021; Mitra et al., 2014). According to traditional ecological theory, keystone species are often defined as those with the most biomass (Kang and Fryxell, 1992; Sergeeva et al., 2018). The combination of CON, CCM and ELA allowed predicting keystone species, i.e. ASVs with strongest effects on the interaction network. We found both highly abundant (for example Fragilariopsis or Pseudo-nitzschia diatoms) as well as low-abundant keystone ASVs, suggesting that both common and rare members contribute to ecosystem stability CCM revealed that by far not all co-occurring ASVs actually influence each other (Fig. 3). A striking example is between clusters 06TS and 05LW, which were closely connected in the CON but not in the CCM network(Fig. 3). This co-occurrence without apparent causal connections could be explained by unique environmental conditions shaping both of these clusters, such as polar water influx. Even more pronounced is the separation of cluster 03LW, mainly heterotrophs, and 08TS, mainly phototrophs, which are tightly connected by co-occurrence but show not a single causal link in the CCM network. The organisms in these two clusters are primarily influenced by environmental parameters, particularly light. Additionally, these photosynthetic and heterotrophic organisms are sometimes preyed upon by the same predators (Zhao et al., 2022) such as Syndiniales. This explains the simultaneous occurrence and similar seasonality of these taxa but indicates that they do not have a direct influence on each other. The lack of causal influence during the transition from polar night to day is clearly visible in the CCM network (see Fig. 3). We interpret this gap between winter and spring clusters as a ‘winter reset’ (Supplementary - Results Figure S11). This phase is characterized by the predominance of Syndiniales and Dinophyceae. With the emergence of light, a new period of primary production begins, shaped by the prevailing environmental conditions. The ambient environmental conditions then determine which species will subsequently prevail. By reflecting causal interactions between species, the CCM network even stronger reflects the cyclic microbiome structure than the co-occurrence network. The cycle begins with photoautotrophs (cluster 08TS) in early spring and ends with the hetero- and mixotrophs (cluster 01TA) in late autumn. As light intensity decreases, mixotrophs become more prevalent than photoautotrophs, leading to a shift towards a heterotrophic lifestyle and a transition from carbon fixation to consumption. This transition into a low light period is characterized by parasitic species, suggesting an “eat and be eaten” scenario. The causal links from fall to winter are significantly less than between other seasons (except winter to spring) Figure 3 B. All of these causal links are related to Syndiniales, which can be explained by their parasitic lifestyle, foraging upon mixotrophic species that are active during the transition autumn phase.

We compared our approach with two previous studies Cross Convergence Mapping (Ushio, 2022; Fujita et al., 2023). While our approach focuses on the specific interactions within and between clusters, Ushio et al. provides a more general framework for predicting community diversity based on interaction capacity, temperature and abundance. The emphasis on mechanistic explanations for observed ecological patterns distinguishes the two approaches. Our methodology provides a comprehensive understanding of keystone species in a specific context, while Ushio’s study provides broader insights into the factors influencing community diversity in different ecosystems. Both studies use similar techniques such as correlation and CCM (Ushio, 2022). Fujita’s study used controlled experiments with six isolated community replicates, subjected to diverse treatments over 110 days. Regarding Takens’ Theorem and Convergence Cross Mapping, Fujita et al. used Simplex projection to forecast population size (Fujita et al., 2023), while our study utilized pairwise CCM on ASV time series signals within clusters to predict keystone species.

The reconstructed landscape for autumn cluster 01TA shows that autumn communities are highly stable. However, for the winter cluster 03LW, the energy values of observed communities lack clear separation, making the situation less straightforward. Communities of the winter clusters which are still present in summer tend to display high energy values, indicating instability in winter conditions. The spring cluster 08TS shows an even more notable trend, indicating that community structures lack stability and exhibit high plasticity under spring conditions (Supplementary - Results Figure S12).

Keystone species represent the ecological roles played by network members in primary production, consumption, and parasitic interactions. During the beginning of autumn, Chlorophyta emerges as a keystone species, indicating a shift in primary production from Ochrophyta to Chlorophyta. This shift may be explained by the preference of Chlorophyta for colder temperatures and a better adaptation to nutrient limitation.

Energy landscape analysis was used to assess the stability of observed communities when exposed to environmental conditions different from their typical settings. Our results showed that communities maintained stable configurations during typical summer (for communities of the summer clusters) or winter (for winter clusters) days, indicating their adaptability to seasonal variations. However, significant changes in energy values occurred when communities were assessed under opposing conditions, suggesting diverse responses to environmental shifts. Subsequent simulations of typical Arctic conditions revealed interesting patterns within the microbial communities. The summer cluster communities showed a clearly decreased stability under Arctic conditions, in contrast to the winter cluster communities, which displayed a tendency towards increased stability. This suggests that winter communities may have a greater capacity to adapt to colder Arctic conditions compared to their summer counterparts (Supplementary - Results Figure S15). The Arctic projection notably harbored a higher number of unique species, indicating varied responses to environmental changes. Notably, the absence of shared keystone species among these different datasets in the summer cluster 10HS suggests a lower robustness of Atlantic ASVs to environmental shifts, with keystone species candidates exhibiting variability. Under Atlantic conditions, the closeness centrality of the summer cluster 10HS increased more compared to Arctic conditions. In the winter cluster community 03LW, closeness centrality is very similar in both projections (Supplementary - Results Figure S15).

The results presented in this study not only have practical implications for ecosystem management by improving our understanding and ability to predict change in complex ecological systems but also provide systematic and mechanistic insights into the mechanisms responsible for shaping and maintaining spatiotemporal heterogeneity in ecosystem composition (Suzuki et al., 2020).

## Supporting information

Supplementary Material - Results

## AUTHOR CONTRIBUTION

WJvA, CB, MW and KM are responsible for the sampling design. KM contributed 18S metabarcoding data. MW processed raw metabarcoding reads into ASVs. WJvA contributed oceanographic data. EO devised the project, the main conceptual ideas and study outline. EO and RMK designed the model and the computational framework and analyzed the data. EO, RMK, OP and KM wrote the initial draft. RMK carried out the implementation. EO, RMK and OP interpreted the data and conceptualized the manuscript. OE and OP advised data evaluation and data interpretation and revised and finalized the manuscript. All authors contributed to the final manuscript, the scientific interpretation and the discussion of results.

## ACKNOWLEDGEMENTS

Computational infrastructure and support were provided by the Centre for Information and Media Technology at Heinrich Heine University Düsseldorf. Thanks to Kenta Suzuki for supporting us in adapting rELA to our needs. Thanks to Taylor Priest for providing the PAR data. We thank Theresa Hargesheimer, Jana Bäger, Jakob Barz, Anja Batzke and Daniel Scholz for the RAS support. Moreover we thank the captains and crews of RV Polarstern for excellent support at sea, and the chief scientists for leading the various expeditions conducted for this study. Ship time for RV Polarstern was provided under grants AWI PS99 00, AWI PS100 01, AWI PS107 05, AWI PS114 01, AWI PS121 01 of RV Polarstern. The work was supported by the infrastructure project FRAM (Frontiers in Arctic Marine Monitoring) funded by the Helmholtz Association. Our special thanks go to Stefan Neuhaus for bioinformatic support, Kerstin Korte and Swantje Ziemann for excellent technical support in the laboratory as well as Martina Löbl for coordination of the FRAM-project.

## SUPPLEMENTARY – METHODS

## DATA AVAILABILITY STATEMENT

The python codes will be available at https://github.com/rakro101/otter.git after acceptance. Raw metabarcoding reads will be accessible under [ENA](https://www.ebi.ac.uk/ena/browser/view/PRJEB43905).

