## Supplementary Material - Methods for "Beyond blooms: A novel time series analysis framework predicts seasonal keystone species and sheds light on Arctic pelagic ecosystem stability"

### 1 DATA PREPROCESSING

#### 1.1 DNA-extraction and Illumina amplicon-sequencing of 18S rRNA genes

Isolation of genomic DNA was carried out using the PowerWater kit (Qiagen, Germany) following the manufacturer's protocol. Obtained DNA was quantified using Quantus (Promega, USA) and stored at -20 °C. 18S rRNA gene fragments from the hypervariable V4 region were amplified by polymerase chain reaction (PCR) with primers 528iF (GCGGTAATTCCAGCTCCAA) and 926iR (ACTTTCGTTCTTGATYRR), illuminaNextV4F (TCGTCGGCA GCGTCAGATGTGTATAAGAGACAGGCGGTAATTCCAGCTCC) and illuminaNextV4R (GTCTCGTGGGCTCGGAGATGTGTATAAGAGACAGGGCAAATGCTTTCGC) (Metfies et al., 2020). All PCRs had a final volume of 50 µL and contained 0.02 U Phusion Polymerase (Thermo Fisher, Germany), the 10-fold polymerase buffer according to manufacturer's specification, 0.8 mM each dNTP (Eppendorf, Germany), 0.2 µM $L^{-1}$ of each primer, and 1 µL of template DNA. PCR amplification was performed in a thermal cycler (Eppendorf, Germany) with an initial denaturation (94 °C,

2 min) followed by 35 cycles of denaturation (94 °C, 20 sec), annealing (58 °C, 30 sec), and extension (68 °C, 30 sec) with a single final extension (68 °C, 10 min). The PCR products were purified from an agarose gel 1% [w/v] with the NucleoSpin Gel Kit (Macherey-Nagel, Germany) and Mini Elute PCR Purification kit (Qiagen, Germany). Subsequently, DNA concentrations were determined using a Quantus Fluorometer (Promega, USA). Prior to library preparation, DNA fragments were diluted with TE buffer to a concentration of 0.2 ng µL-1. Libraries were prepared according to the 16S Metagenomic Sequencing Library Preparation protocol, and sequenced using MiSeq (Illumina, USA) in 2x300 paired-end runs. Sequence data are available under ENA BioProjects PRJEB43889 and PRJEB43890, with accession numbers of individual fastq files shown in Supplementary Table X.

#### 1.2 Sequence analysis

After primer removal using cutadapt (Martin, 2011) reads were processed into amplicon sequence variants (ASVs) using DADA2 v1.14.1 (Callahan et al., 2016), as described in Wietz el al (2021). Briefly, reads were trimmed based on quality profiles, with filtering settings truncLen=c(250,200), maxN=0, minQ=2, maxEE=$c(3, 3)$ and trunc$Q = 0$. Followed by merging (minOverlap= $20$) and chimera removal, reads were taxonomically classified using PR2 v4.12. The herein reported data has been processed in the scope of autonomous eDNA biodiversity analyses within the FRAM Observatory, as described under `https://github.com/matthiaswietz/FRAM-RAS_eDNA`.

#### 1.3 Hellinger Transformation

In the matrix $\mathbf{M}$, the columns represent observed species, and the rows represent different samples. The entries are the raw abundance of the species at given sample. The Hellinger transformation was applied to facilitate data normalization and comparison of ecological data, reducing the impact of differences in the scale of abundance values across samples. The normalisation is applied column-wise to the raw abundance data as follows:

1. Calculate the square root of each element:

$$\mathbf{S} = \sqrt{\mathbf{M}}$$

2. Calculate the L2 norm (Euclidean norm) of each column in $\mathbf{S}$:

$$\mathbf{R} = \|\mathbf{S}\|_2 \text{ (column-wise)}$$

3. Normalize each column by dividing it by its L2 norm:

$$\mathbf{H} = \frac{\mathbf{S}}{\mathbf{R}}$$

### 2 CO-OCCURRENCE NETWORK

#### 2.1 Fourier Transformation of Time Series

Oscillation signals were computed for each ASV in yearly datasets using discrete Fourier transformation (first 14 Fourier Components) of normalized abundances. This process entailed the conversion of temporal variations within each ASV's abundance into the frequency domain, enabling the capture of cyclic patterns indicative of potential ecological interactions (Machné et al., 2017).

#### 2.2 Calculation of Pearson Correlation

The correlation between two abundance time series $Y$ and $\hat{Y}|M_x$ can be calculated using the formula:

$$\text{corr}(Y, \hat{Y}) = \frac{\text{cov}(Y, \hat{Y})}{\sqrt{\text{var}(Y) \cdot \text{var}(\hat{Y})}}$$

where $\text{cov}(Y, \hat{Y})$ is the covariance between $Y$ and $\hat{Y}$, $\text{var}(Y)$ is the variance of $Y$, and $\text{var}(\hat{Y})$ is the variance of $\hat{Y}$.

### 3 CONVERGENT CROSS MAPPING

#### 3.1 Attractors

An attractor represents a group of states towards which a system naturally converges over time. When these attractors arranged in a complex, often non-linear or curved manner, they are called ”manifolds.” A useful analogy is to imagine attractors as a way of summarising our system, similar to an embedding layer in neural network architectures (Borja et al., 2020).

#### 3.2 Takens' Embedding Theorem and Shadow Manifolds

Emerging from the term ”shadow,” shadow manifolds serve as ”projections” of the genuine system manifold onto a specific variable $X$. Takens' theorem provides insights into constructing a shadow manifold $M_x$ that mirrors the true manifold $M_s$ by utilizing lagged (historical) values of $X$. The points within the shadow manifold $M_x$ establish a one-to-one correspondence with the points within the actual (unobservable) manifold $M_s$ (Uzal et al., 2011).

#### 3.3 Cross Mapping

Given points on the manifold of one variable $M_y$, we seek corresponding points on the manifold $M_x$, i.e., points at the same time $t$. If variable $X$ has an effect on the response of variable $Y$, we infer that information about $X$ is encoded in $Y$. Hence, predictions of $X$ values can be made using information from $Y$. Notably, this concept differs from Granger causality, which suggests that $X$ causes $Y$ if predicting $Y$ is improved by knowing $X$. Shadow manifolds are employed to navigate the challenge of unknown manifolds. While the true system manifold is often elusive, Takens' theorem offers a framework for constructing shadow manifolds $M_x$ and $M_y$, which can be cross-mapped with a one-to-one correspondence to the true system manifold (Takens, 1981). Using an analogy, manifolds can be considered condensed summaries of a system's behaviour. In this analogy, $M_x$ and $M_y$ act as abstract representations of variables $X$ and $Y$, respectively. Portions of $X$ are effectively embedded within $Y$ when $X$ influences $Y$. We can, as a result, use $M_x$ to make predictions about $Y$, which we denote as $\hat{Y}|M_x$. The accuracy of this prediction, typically measured by metrics such as Mean Absolute Error (MAE), Mean Squared Error (MSE), or correlation, serves as a measure for inferring causal relationships. More generally, the effectiveness of the predictions made using $M_x$ for $Y$ is used as a way of assessing potential causal relationships between the variables $X$ and $Y$ (Borja et al., 2020; Sugihara et al., 2012).

#### 3.4 Convergence

Convergent Cross Mapping (CCM) centres on variables with causal links. The central concept is that the increase in observation period or data collection enhances the ability to forecast one variable based on another. The attractor example, previously mentioned, supports this notion. As time progresses, the attractor becomes denser, as the system fills gaps in the manifold. Essentially, more well-defined manifolds suggest that variables with causal connections are likely to exhibit improved accuracy of predictions, such as $\hat{Y}|M_x$. On the contrary, if two variables lack a causal link, improving their manifolds might not lead to enhanced predictions. In the CCM framework, both cross-mapping and convergence are vital prerequisites to explore causal relationships between variables (Borja et al., 2020; Sugihara et al., 2012).

We utilize the CCM Algorithm but substitute the Pearson correlation with the normalized mutual information score (Javier, 2021). Here, we choose to use mutual information because it is directed, whereas the Pearson correlation is not direct, i.e. $p(x, y) = p(y, x)$, but $nmi(x, y)! = nmi(y, x)$, where $x$ and $y$ are two time series signals. Hence, our dataset has under 100 samples over the four years, we consider it more as discrete data set and use the normalized mutual information score instead of Pearson correlation, that assumes continuous variables (Ross, 2014; Liu, 2019). In addition, Pearson correlation has the assumption of linear relations ships, where mutual information can deal with non-linear relations (Ross, 2014; Liu, 2019). While a 'p-value' test for strict independence may decrease significantly when applied to a large dataset with even slightly related variables, Mutual Information (MI) will converge to a measure of their relatedness with a high degree of accuracy (Ross, 2014).

#### 3.5 Algorithm Description

Given two time series $X = \{X_1, X_2, \ldots, X_L\}$ and $Y = \{Y_1, Y_2, \ldots, Y_L\}$, where $L$ is the time series length:

1. Calculate lagged-coordinate vectors using the following formula:

$$x_t = \langle X_t, X_{t-\tau}, X_{t-2\tau}, \ldots, X_{t-(E-1)\tau} \rangle$$
$$\text{for } t \in [1 + (E-1)\tau, L]$$

   where $E$ is the embedding dimension, and $\tau$ is the lag step.
2. Define the "shadow (attractor) manifold" $M_x = \{x_t \text{ for each } t \in [1 + (E-1)\tau, L]\}$.
3. Identify the $E + 1$ nearest neighbor vectors from the selected vector $x_t$ within $M_x$.
4. Denote the time indices of the $E + 1$ nearest neighbors of $x_t$ as $t_1, \ldots, t_{E+1}$, which will be used to identify corresponding points in $Y$.
5. Define the model predicting $Y$ given $M_x$ using the formula:

$$\hat{Y}|M_x = \sum_{i=1}^{E+1} w_i Y_{t_i}$$

   where $w_i$ represents the weight multiplied by $Y_{t_i}$:

$$w_i = \frac{u_i}{\sum_{j=1}^{E+1} u_j}, \quad \text{where } j = 1 \ldots E+1$$

   and

$$u_i = \exp\left[-\frac{d(x_t, x_{t_i})}{d(x_t, x_{t_1})}\right]$$

Here, $d(x_s, x_t)$ represents the Euclidean distance, and the division by $d(x_t, x_{t_1})$ scales the distances as multiples of the distance to the closest point.

6. Calculate the normalized mutual information score between $Y$ and $\hat{Y}|M_x$ using the formula:

$$\text{Normalized Mutual Information} = \frac{I(Y;\hat{Y}|M_x)}{\sqrt{H(Y)\cdot H(\hat{Y}|M_x)}}$$

where $I(Y;\hat{Y}|M_x)$ is the mutual information between $Y$ and $\hat{Y}|M_x$, and $H(Y)$ and $H(\hat{Y}|M_x)$ are the entropies of $Y$ and $\hat{Y}|M_x$, respectively.

7. To assess the statistical significance of the normalized mutual information score, the permutation test can be employed. The basic idea is to repeatedly shuffle the values of $Y$ while keeping the mapping between $Y$ and $\hat{Y}|M_x$ intact. For each permutation, recalculate the normalized mutual information score. This generates a null distribution of scores under the assumption of no true relationship between $Y$ and $\hat{Y}|M_x$.
8. The p-value is then computed as the proportion of permuted scores that are greater than or equal to the observed normalized mutual information score. In mathematical terms, if $N$ is the number of permutations and $s_i$ is the normalized mutual information score for the $i$-th permutation, the p-value can be calculated as:

$$\text{p-value} = \frac{1}{N}\sum_{i=1}^{N}\mathbb{I}(s_i \geq \text{observed score})$$

where $\mathbb{I}$ is the indicator function.

9. The obtained p-value helps determine whether the observed normalized mutual information score is statistically significant, indicating a meaningful relationship between $Y$ and $\hat{Y}|M_x$.

#### 3.6 Application on our data

Convergent Cross Mapping (CCM) is a method for analysing potential causal relationships between variables in time series data. It attempts to determine whether changes in one component are associated with changes in another component. In a first step we built a CCM-Network from all pairwise combination comprising all species abundance data. Secondly we extracted the in- and outgoing edges between the nodes that are connected in the co-occurrence network.

We used the implementation of Normalized Mutual Information (NMI) from `https://github.com/polsys/ennemi` by Petri Laarne and the Convergent Cross Mapping by Implementation from Javier, Prince`https://github.com/PrinceJavier/causal_ccm` (Javier et al., 2022) to measure the strength of the causal relationship.

**Remark**: We also added normalised mutual information to Sugihara's cpp ($C^{++}$) implementation (Sugihara et al., 2012), but chose to include Javier's pure Python implementation for our experiments. Thus CCM is only one module of our approach, Sugihara's PyEDM package can easily be used instead of Javier's implementation (Javier et al., 2022).

For the Convergent Cross Mapping (CCM) algorithm, we configured the following parameters:
**Time Lag** $(\tau) = 1$,
**Embedding Dimension** $(d) = 2$,

**Time Period/Duration** $(L) = 96$**.**
The time period was determined by the number of samples.

#### 3.7 Modified CCM can detect predator-prey relationships

To demonstrate the effectiveness of the modified CCM, we analyzed time series data from a traditional predator-prey experimental system that was initially researched in the 1920s by Gause and subsequently refined by Veilleux. (Veilleux, 1979; Sugihara et al., 2012). The system involves Didinium (predator) and Paramecium (prey). We sourced the data exactly as provided, which can be found at `https://robjhyndman.com/tsdldata/data/veilleux.dat`, and utilized only the most recent 62 data points, as recommended by (Sugihara et al., 2012).

#### 3.8 CCM connectivity statistics

To validate the robustness of our findings, we calculated a non-parametric P-values using 1000 random permutations of the abundance matrix. The significance of edge weights derived from the CCM was assessed by comparing them to those obtained from the random network. Here the p-value denotes the ratio of weights that are higher in CCM compared to random, thus enabling a bijective mapping from normalized mutual information (edge weights) to p-value (see Supplementary Material Results Figure GQR3). Similar non-parametric p-value definition are used for CO-Occurrence Networks (Ma et al., 2016).

In the next step, we investigated whether the current co-occurrence edges exhibit greater NMI values in comparison to random and hypothetical edges. For this purpose, we established three separate networks: (i) integrating edges from the co-occurrence network, (ii) incorporating unconnected nodes from the co-occurrence network, and (iii) utilizing randomly generated data.

#### 3.9 Cluster aggregation into a single node

To enhance the elucidation of inter-cluster interactions, we employ an aggregation approach within the context of the present methodological framework. Specifically, we aggregate all edges originating from nodes situated within e.g. Cluster A and terminating in Cluster B, utilizing the arithmetic mean of normalized mutual information (NMI) as the amalgamation criterion:

$$\text{Mean NMI (A}\rightarrow\text{B)} = \frac{1}{N}\sum_{i=1}^{N}\text{NMI}(A_i, B_i) \tag{S1}$$

Where: - $N$ represents the total number of edges between Cluster A and Cluster B. - NMI$(A_i, B_i)$ denotes the normalized mutual information between node $A_i$ in Cluster A and node $B_i$ in Cluster B.

This procedure effectively condenses the quantity of items to be depicted within the node cloud, ultimately resulting in the representation of clusters by a singular, composite node.

#### 3.10 Definition: Network distance

We employed Fourier-transformed abundance values time series signals to generate temporal profiles for eukaryotic ASVs. UMAP was applied to the temporal profiles, resulting in a three-dimensional embedding space (McInnes et al., 2018). Within this space, centroids were calculated for each cluster (Supplementary Material Results Figure S2,S3,S4). The network distance between clusters was quantified as the Euclidean distance between their respective centroids. Subsequently, a distance matrix was constructed and distances were rounded to integers. The matrix was pruned by only significant connections.

#### 3.11 Closeness Centrality and Betweenness Centrality

To compute closeness centrality and betweenness centrality, we use 1-NMI as the distance/edge weight, where NMI is the directed normalised mutual information between nodes (Freeman et al., 2002).

### 4 ENERGY LANDSCAPE ANALYSIS

The following definitions and algorithms are based on (Suzuki et al., 2021; Fujita et al., 2023).

#### 4.1 Definition of State Space

Formalizing the stability landscape concept yields a definitive definition of the state space within an energy landscape. We denote a community composition, represented as a binary vector of length $S$, where $S$ signifies the total number of species. Within this framework, there exist $2^S$ unique community compositions corresponding to the nodes of the stable state. Specifically, a community composition of the $k$-th sample ($k \in \{0, 1, ..., 2^S - 1\}$) is denoted as $\sigma^{(k)} = (\sigma_1^{(k)}, \sigma_2^{(k)}, ..., \sigma_S^{(k)}$, where $\sigma_i^{(k)} \in \{0, 1\}$ indicates the presence/absence status of the $i$-th species.

To establish links between community compositions, we adopt the assumption that transitions occur incrementally. Consequently, two community compositions are linked if and only if they differ in the presence/absence status of precisely one species. This leads to the formation of a structured network wherein each node is connected to $S$ neighbors.

#### 4.2 Pairwise Maximum Entropy Models and Energy Landscape

We attribute energy values to individual community compositions and establish the potential structure within the state space through the introduction of the extended pairwise maximum entropy model. This model governs the likelihood of observing community composition $\sigma^{(k)}$ under an environmental condition $\epsilon = (\epsilon_1, \epsilon_2, ..., \epsilon_M)$, where $\epsilon_i$ represents continuous values denoting environmental factors like resource availability, pH, temperature, or host organism age. The probability of $\sigma^{(k)}$ occurring in condition $\epsilon$ is given by:

$$P(\sigma^{(k)}|\epsilon) = \frac{e^{-E(\sigma^{(k)},\epsilon)}}{Z}, \tag{S2}$$

with the energy defined as:

$$E(\sigma^{(k)}, \epsilon) = -\sum_{i=1}^{S} h_i \sigma_i^{(k)} - \sum_{i=1}^{S}\sum_{j=1}^{M} g_{ij}\epsilon_i \sigma_j^{(k)} - \sum_{j=1}^{S}\sum_{i=1, i\neq j}^{S} J_{ij}\sigma_i^{(k)}\sigma_j^{(k)}/2, \tag{S3}$$

where $E(\sigma^{(k)}, \epsilon)$ represents the energy of community composition $\sigma^{(k)}$ and

$$Z = \sum_{k=0}^{2^S-1} e^{-E(\sigma^{(k)},\epsilon)}. \tag{S4}$$

It is pertinent to note that the term $E(\sigma^{(k)}, \epsilon)$ is labeled as energy due to its analog in statistical physics (Azaele et al., 2010), although it serves as an exponent in Equation (S2) and indicates the likelihood of observing a community composition within an ecological context. It does not correspond directly to physical energy as used in ecological studies. The parameters in Equation (S3) encompass $h_i$, $J_{ij}(S \times S)$ , and $g_{ij}(M \times S)$, which are components of vector $\mathbf{h} = (h_1, h_2, ..., h_S)$, matrix $\mathbf{J} = (J_{ij})$, and matrix $\mathbf{g} = (g_{ij})$ respectively. Here, $h_i$ signifies the net impact of unobserved environmental factors favoring ($h_i > 0$) or hindering ($h_i < 0$) the presence of species $i$, and $g_{ij}$ represents the influence of the $i$-th observed environmental factor on the occurrence of species $j$. The model captures pairwise relationships, as each species is interconnected with all others through $J_{ij}$.

The extended pairwise maximum entropy model reduces to the pairwise maximum entropy model when the influence of environmental conditions is negligible:

$$E(\sigma^{(k)}) = -\sum_{i=1}^{S} h_i \sigma_i^{(k)} - \sum_{j=1}^{S} \sum_{i=1, i \neq j}^{S} J_{ij} \sigma_i^{(k)} \sigma_j^{(k)} / 2. \tag{S5}$$

see (Azaele et al., 2010; Harris, 2015; Araújo et al., 2011). EEquation (S3) and Equation (S5) account for pairwise relationships, assuming that the information encapsulated in the first two moments sufficiently characterizes higher-order occurrence patterns (Azaele et al., 2010). Although this assumption can be extended to include higher-order terms, it necessitates larger data sets for accurate prediction (Nguyen et al., 2017).

The pairwise maximum entropy models associate energy values with each community composition in the state space. Energy signifies the directionality of transitions between community compositions. For two adjacent nodes, $\sigma^{(k)}$ and $\sigma^{(k_0)}$, if $E(\sigma^{(k)}) > E(\sigma^{(k_0)})$, then the transition from $\sigma^{(k)}$ to $\sigma^{(k_0)}$ is more likely than the reverse. Such local transition rules govern dynamics, and the energy landscape dictates the overall compositional dynamics. This may approximate the stability landscape.

Assuming a data set comprising community compositions of $N$ samples, denoted as matrix $\mathbf{X} = (x_1, x_2, ..., x_N)(S \times N)$ where $x_i \in \sigma_1, \sigma_2, ..., \sigma_{2^S}$, and optionally, environmental conditions $\mathbf{Y} = (y_1, y_2, ..., y_N)(M \times N)$ where $y_i = \epsilon^i = (\epsilon_1^i, \epsilon_2^i, ..., \epsilon_M^i)$ denotes the environmental factors of the $i$-th sample, we can obtain maximum likelihood estimates for $h$, $J$, and $g$. These estimates can be acquired through gradient descent for the pairwise maximum entropy model (Equation (S5)) or stochastic approximation for the extended pairwise maximum entropy model (Equation (S3)), as described in the Appendix.

Given that these equations emitted from the maximum entropy principle, the probability distribution of community compositions under these parameters represents an unbiased estimate based on observational data (Jaynes, 1982; Harte and Newman, 2014). In essence, this probability distribution conforms to the observational data constraints while maximizing remaining uncertainty, making it a well-established assumption for estimating the occurrence probability of unobserved community compositions (Elith* et al., 2006; Shipley et al., 2006; Parisien and Moritz, 2009; Franklin, 2010; Staniczenko et al., 2017; Clark et al., 2018).

Energy landscape analysis delves into the topological and connectivity attributes of an energy landscape. In the following sections, we elucidate the identification and significance of key components within this analysis and elaborate on how they contribute to our understanding of stability landscape structure. The term ”node” is employed in a similar context as that of a community composition.

### 4.3 Energy Minima

We designate energy minima within an energy landscape as stable states in the corresponding stability landscape- These local minima exhibit the lowest energy values in comparison to their neighboring nodes, thus representing endpoints in scenarios where assembly processes are entirely deterministic (i.e., transitions of community compositions consistently descend the energy landscape). The existence of multistability manifests as the presence of multiple energy minima within a given energy landscape. Given that an energy minimum denotes a node with lower energy than all adjacent nodes, we assessed whether each of the $2^S$ nodes qualified as local minima. This concept, although briefly mentioned in (Azaele et al., 2010)), resonates with our analysis of marine community distributions.

### 4.4 Basin of Attraction

The basin of attraction encompasses the collection of community compositions that converge to a distinct stable state under the assumption of completely deterministic assembly processes. Identification of the basin of attraction is grounded in the stable state to which a given composition converges. Our approach to determining the basin of attraction employs the steepest descent method, as outlined below. Commencing with the selection of a node $i$ within the energy landscape, if the chosen node is not a local minimum, we transition to the node possessing the lowest energy value among its adjacent counterparts. Iteratively, traversing downhill through this process continues until a local minimum is reached. The initial node $i$ is then attributed to the basin associated with the identified local minimum. This procedure is repeated for each of the $2^S$ nodes, excluding those corresponding to local minima.

### 4.5 Disconnectivity Graph

A disconnectivity graph, serves as a concise representation of the hierarchical relationships existing among stable states within an energy landscape. The terminal leaves of this tree correspond to the stable states, their vertical positions reflect their respective energy values, and the branches depict the energy barriers that separate these stable states. The junction where community compositions link two branches signifies a tipping point. This tipping point represents the nadir of the ridge situated between two basins. When contemplating the transition from a stable state $\sigma_A$ to another stable state $\sigma_B$ ($\sigma_A \to \sigma_B$), the energy barrier's height is computed as the difference between the energy at the tipping point and that of $\sigma_A$. Similarly, the energy barrier's height for the transition $\sigma_B \to \sigma_A$ is calculated as the difference between the energy at the tipping point and that of $\sigma_B$. Consequently, transitions between two stable states typically exhibit asymmetrical directionality, and transitions involving smaller energy barriers tend to occur more frequently than their counterparts. Determining tipping points involves examining the connectivity of the energy landscape while altering an energy threshold value through the following steps:

1. Set an energy threshold value, denoted as $E_{\text{th}}$, equal to the energy of the node with the second-highest energy value among all community compositions.
2. Remove the node (along with its associated links) having the highest energy value such that the node with energy $E_{\text{th}}$ attains the highest energy state.
3. Check for connectivity between each pair of stable states within the modified network.
4. Decrease $E_{\text{th}}$ back to the second-highest energy value and repeat the process until all local minima are isolated.
5. For each pair of stable states, record a community composition with the lowest energy value below which the two stable states become disconnected.

Any implausible stable states are pruned, specifically those with the shallowest energy barrier 20% lower than the highest energy barrier across the entire energy landscape.

#### 4.6 Stable State Diagram

The stable state diagram, elucidates the energies of stable states and tipping points featured within disconnectivity graphs across distinct environmental conditions. This diagram provides insights into the alterations an energy landscape undergoes in response to shifts in environmental gradients. Each stable state (and its corresponding tipping point) is represented by a line segment signifying the range of environmental conditions within which it is identifiable. Typically, stable states are depicted as solid lines, whereas tipping points are represented by dashed lines.

#### 4.7 Application of Energy Landscape Analysis

In this study, we analyzed the Energy Landscape by utilizing the following parameters. For each cluster, we only considered the Top 100 ASVs: **Relative Abundance Threshold:** For non-low light clusters, a threshold of $2\%$ was applied, while for low light clusters, we used a threshold of $1\%$. **Occurrence Threshold (Lower):** We set the occurrence threshold (lower) to $1\%$. **Occurrence Threshold (Upper):** The occurrence threshold (upper) was defined as $99\%$. **Stable State Selection:** A pruning threshold of $5\%$ was used to select stable states. **Computational Settings:** All calculations were performed using 20 threads, and we ran 512 replicates for the Relative Setup. **Computation Threshold:** We specified a $q_{\text{th}}$ value of $10 \times 10^{-5}$ as the threshold for stopping computations. For more detailed information, please checkout our GitHub repository.

To improve the visual clarity of the rELA 3D plots, we filtered out community composition data points where the MDS1 and MDS2 coordinate exceeded the mean plus/ minus twice the standard deviation of the community composition within the plot. This step was taken to enhance visualization and eliminate outliers.

#### 4.8 Estimating stability of communities in different environments

To calculate the energy landscape and the stable states for the communities, we use the following three interaction parameter: i) $\mathbf{J}$ which denotes the interaction between species ii) $\mathbf{G}$, which represents the interaction between species and environment, and iii) the vector $h$ which contain the effects of unobserved environmental factors.The median conditions for the time period of interest (either Arctic Summer or Arctic Winter) represent the environmental data of a specific season. This vector is employed in the calculations of the Energy Landscape Algorithm (ELA). When projecting these calculations to the Arctic, we make assumptions based on the environmental vector given in Table S0. Subsequently, we aim to integrate the predicted stable community species into the time series data of polar regions and assess the impact on the overall species abundance. We simulated only winter and summer, specifying clear external conditions like no light and high polar water influence. The transition phases, like spring and autumn, are too variable in environmental conditions to set fixed parameters.

### SUPPLEMENTARY MATERIAL RESULTS

**Table S0.** Environmental Parameters for Arctic Summer and Winter, Atlantic winter and summer. Photosynthetically Active Radiation (PAR). The Arctic (Arc.) Summer and Winter values are set. The Atlantic (Atl.) Summer and Winter values are calculated as the median of the respective seasons of the four years.

| Parameter | Description | Arc. Summer | Arc. Winter | Atl. Summer | Atl. Winter |
|---|---|---|---|---|---|
| MLD | Mixed Layer Depth [m] | 0 | 0 | 0 | 252 |
| PAR | PAR [$\mu$ mol photons $m^-2d^-1$] | 40 | 0 | 27 | 0 |
| Temp | Temperature [°C] | -2 | -2 | 5.81 | 3.82 |
| Salinity | Salinity [PSU] | 34 | 34 | 35 | 34.99 |
| PW_Frac | Polar Water fraction [%] | 100 | 100 | 11 | 11 |
| O2_conc | Oxygen Concentration [$\mu$mol $l^{-1}$] | 350 | 305 | 318 | 303 |
| Depth | Depth [m] | 30 | 40 | 26 | 33 |
