## Supplementary Material - Results for "Beyond blooms: A novel time series analysis framework predicts seasonal keystone species and sheds light on Arctic pelagic ecosystem stability"

### 1 BIO-DIVERSITY

Alpha diversity describes the biodiversity within a specific area or local environment, such as a particular habitat or community. It evaluates the variety and abundance of species within a defined location. Various metrics such as the Simpson Index, Shannon Entropy, and Chao Richness were employed to calculate alpha diversity in our study (Simpson, 1949; Shannon, 1948; Johnson and Angeler, 2014). These metrics offer insights into species richness, which is the count of unique species, and evenness, which measures the distribution of individuals among species in each cluster of ASVs. ASVs are defined by co-occurrence patterns. The Simpson's Index assesses the probability of two individuals chosen at random from a sample belonging to the same species, while the Shannon's Entropy takes into account both species richness and evenness in an ecosystem. Chao1 Richness estimates the quantity of rare species, facilitating a more thorough comprehension of diversity within our specified clusters (Figure S1).

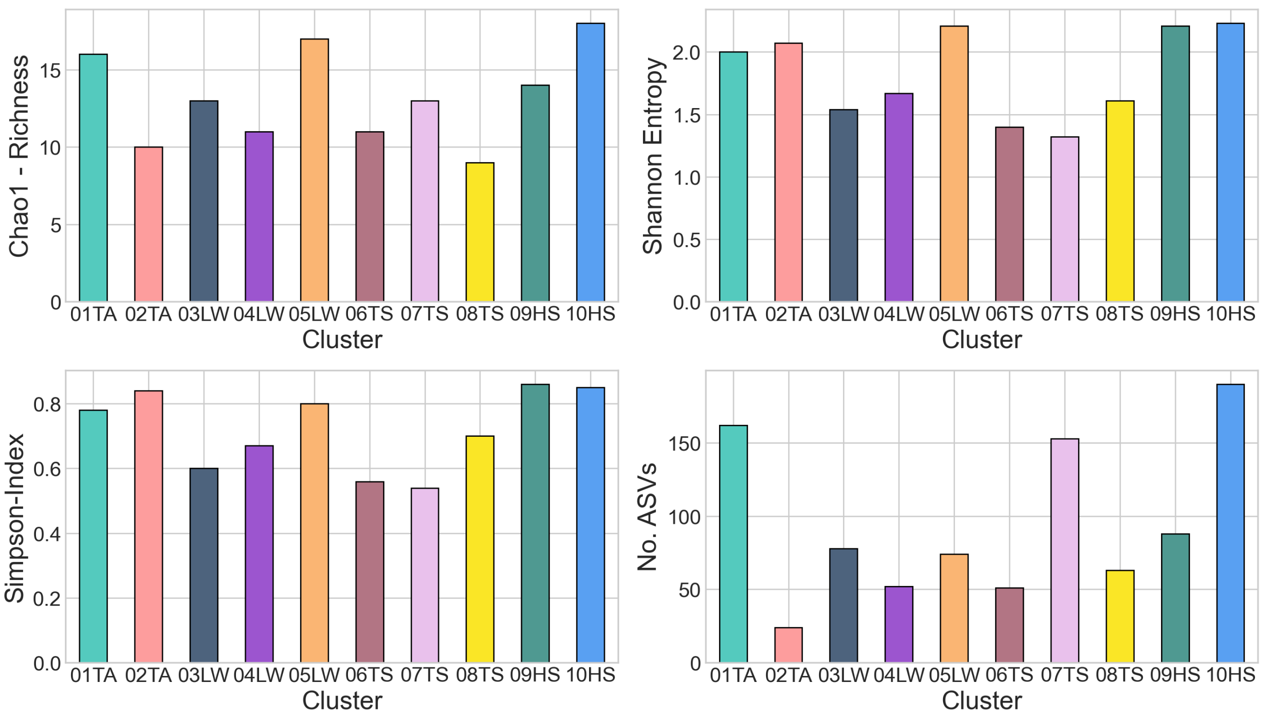


**Figure S1. (Alpha) Bio-Diversity for the ten clusters:** Phylum level biodiversity was measured using Simpson's index, Shannon's entropy and Chao1 richness, and the number for ASV for each cluster.

In contrast, beta diversity highlights variations in the types of species present in different environments or communities, emphasising the diversity differences between these distinct regions, rather than within them. In your study, beta diversity was computed using the Bray-Curtis distance metric (Beals, 1984). This measure assesses the dissimilarity in species composition between different clusters or groups. In this case, it unveils disparities in species composition among co-occurring pattern-based clusters. It presents a quantitative assessment of the variability in species' incidence or density among distinct groups, aiding the comprehension of the degree of similarity or dissimilarity in species composition among different ecological clusters (Figure S2).

All calculations in this subsection was utilized by the python package scikit bio `https://github.com/scikit-bio/scikit-bio` and pandas (`https://github.com/pandas-dev/pandas`) on normalized count data of the taxa level Phylum.

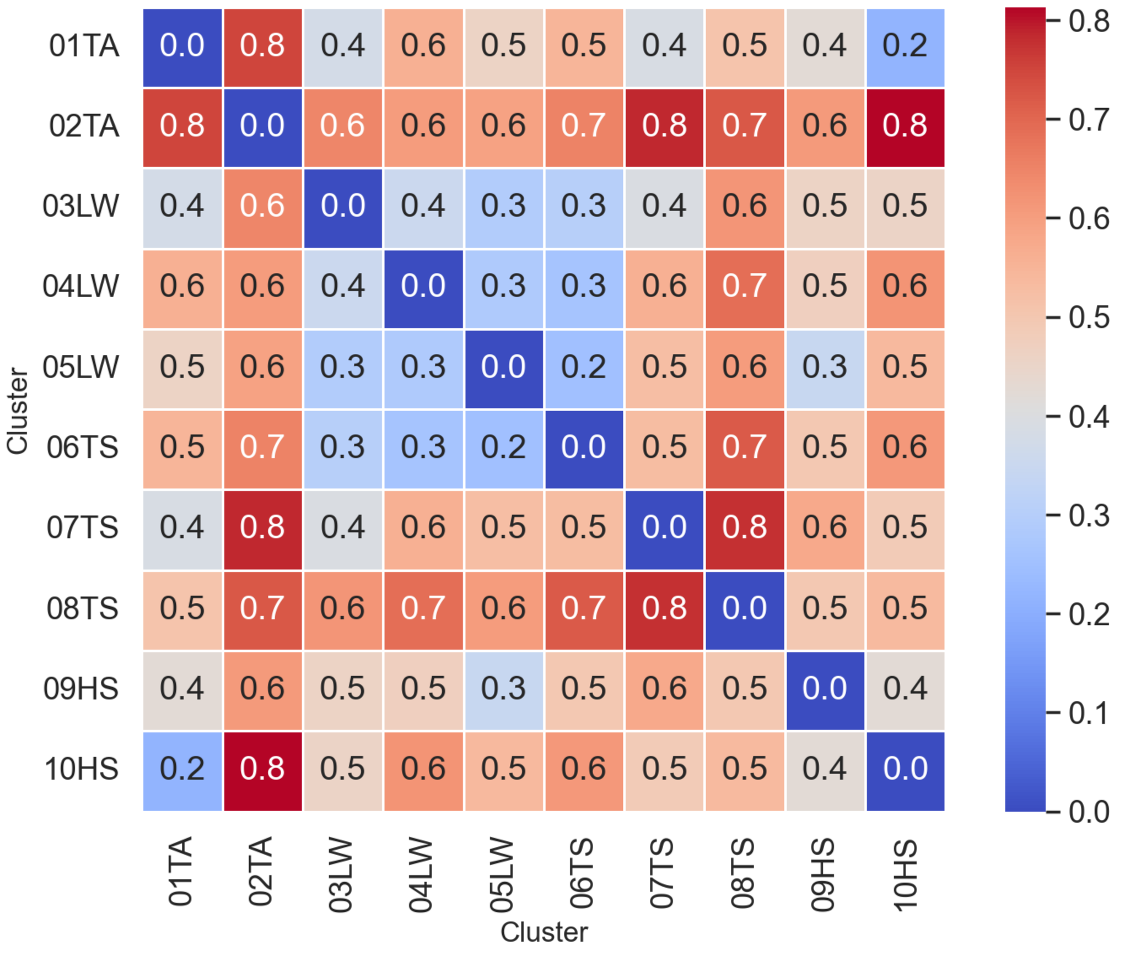


**Figure S2. Pairwise (Beta) Bio-Diversity for the ten clusters:** The beta diversity, measured using Bray-Curtis, was used to determine the biodiversity of each cluster on a Phylum level. The colours indicate the magnitude of the beta diversity coefficient.

### 2 CONVERGENT CROSS MAPPING

In investigating CCM networks, standard methods commonly use techniques such as Pearson correlation to assess the connectivity and dynamics between ASV time series signals within a system, represented in our study by a eukaryotic community over a four-year period. Using Normalised Mutual Information (NMI) to construct a CCM network provides a distinctive method to assess the links between ASV time series signals. NMI works as a metric to compute the amount of information one ASV time series signal contains about another, efficiently quantifying their interdependence by examining the shared information between them. This method differs from the usual Pearson correlation approach, which concentrates on assessing linear associations, comprising the extent and direction of such connections within ASV time series signs. The use of NMI in the CCM framework allows for a refined exploration of the dependencies and information flows within the system, enabling the identification of non-linear relationships that might otherwise be obscured by traditional correlation-based analyses.

#### 2.1 Normalized Mutual Information and p-value definition

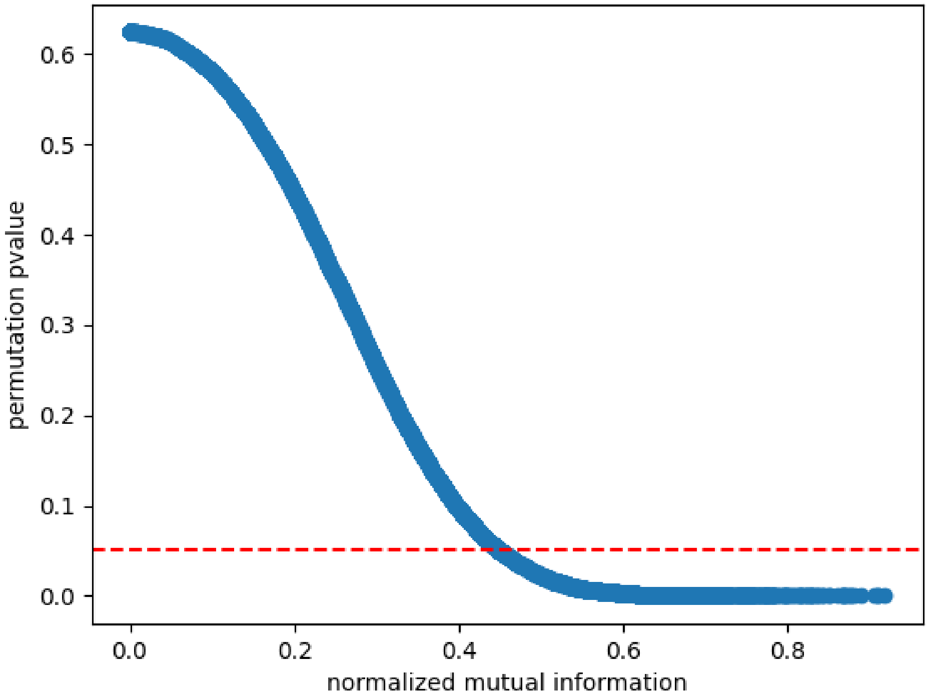


**Figure S3. Significance of NMI Edge Weights.** This plot shows the bijective mapping from normalized mutual infomation and the ($\Psi$) p-value in blue. The red line is the threshold $p < 0.05$.

### 2.2 Network distance calculation

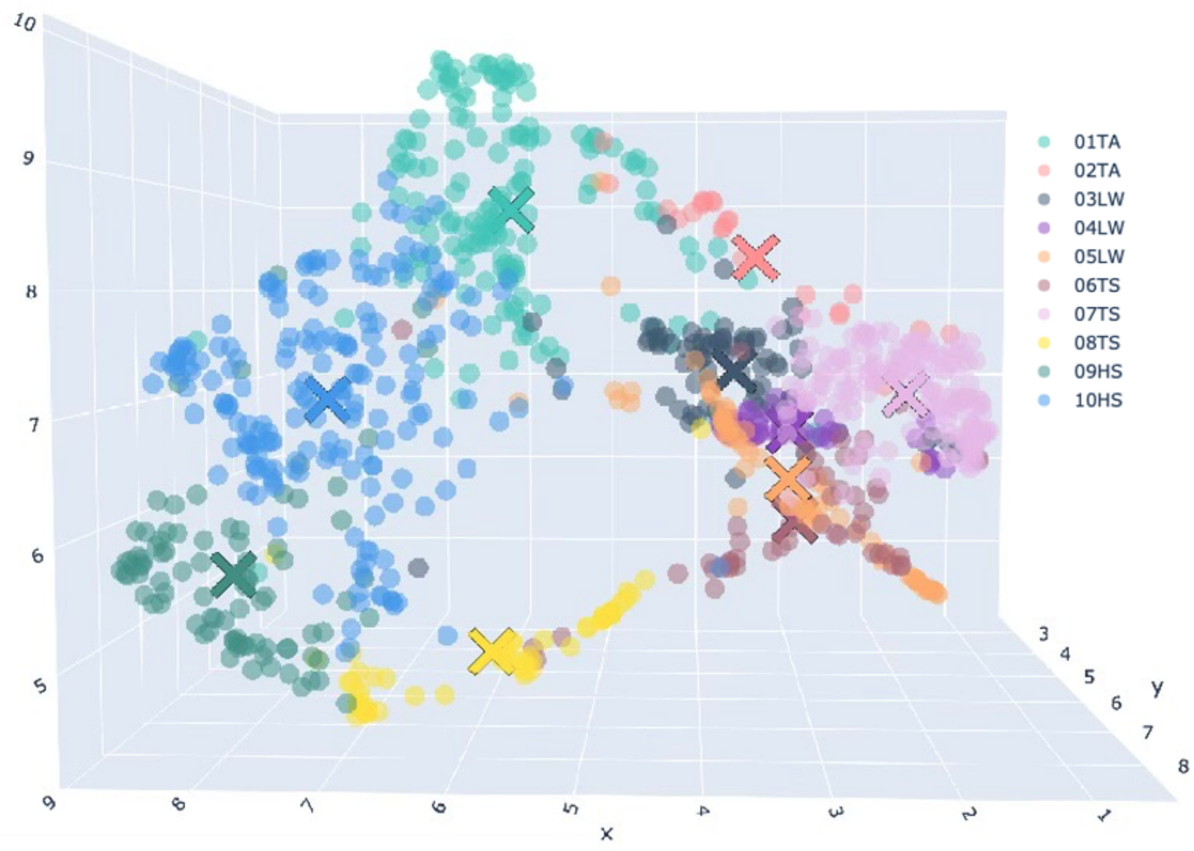


**Figure S4. Umap Projection 3d of temporal profiles:** Projection of the temporal profiles of Eukaryotes. The balls are the ASVS and cross are the cluster centroids. The color indicate the cluster assignment.

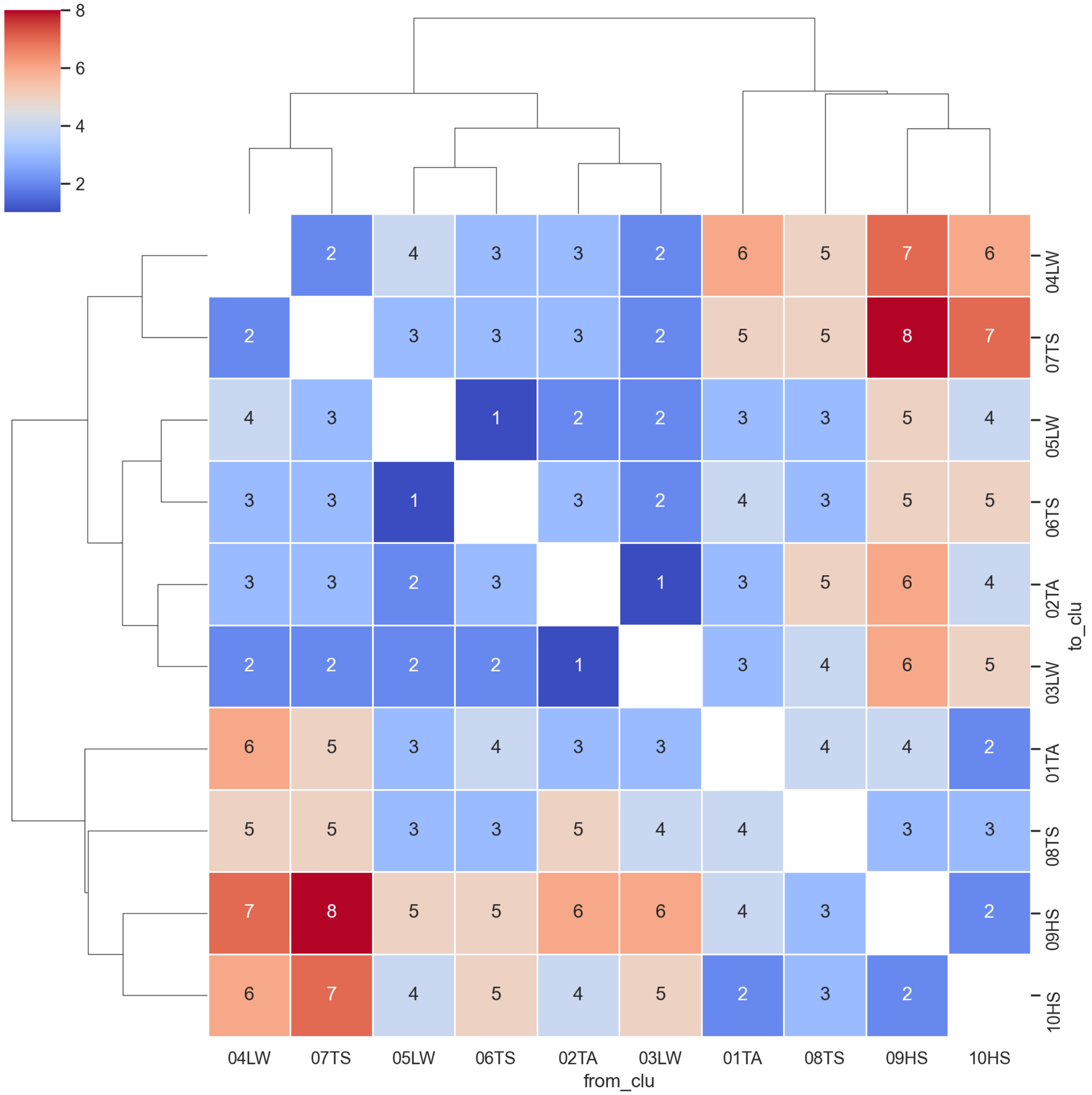


**Figure S5. Distance from Umap Projection 3d** Euklidian distance of the 3d Projection of the temporal profiles of Eukaryotes based on the cluster centroids.

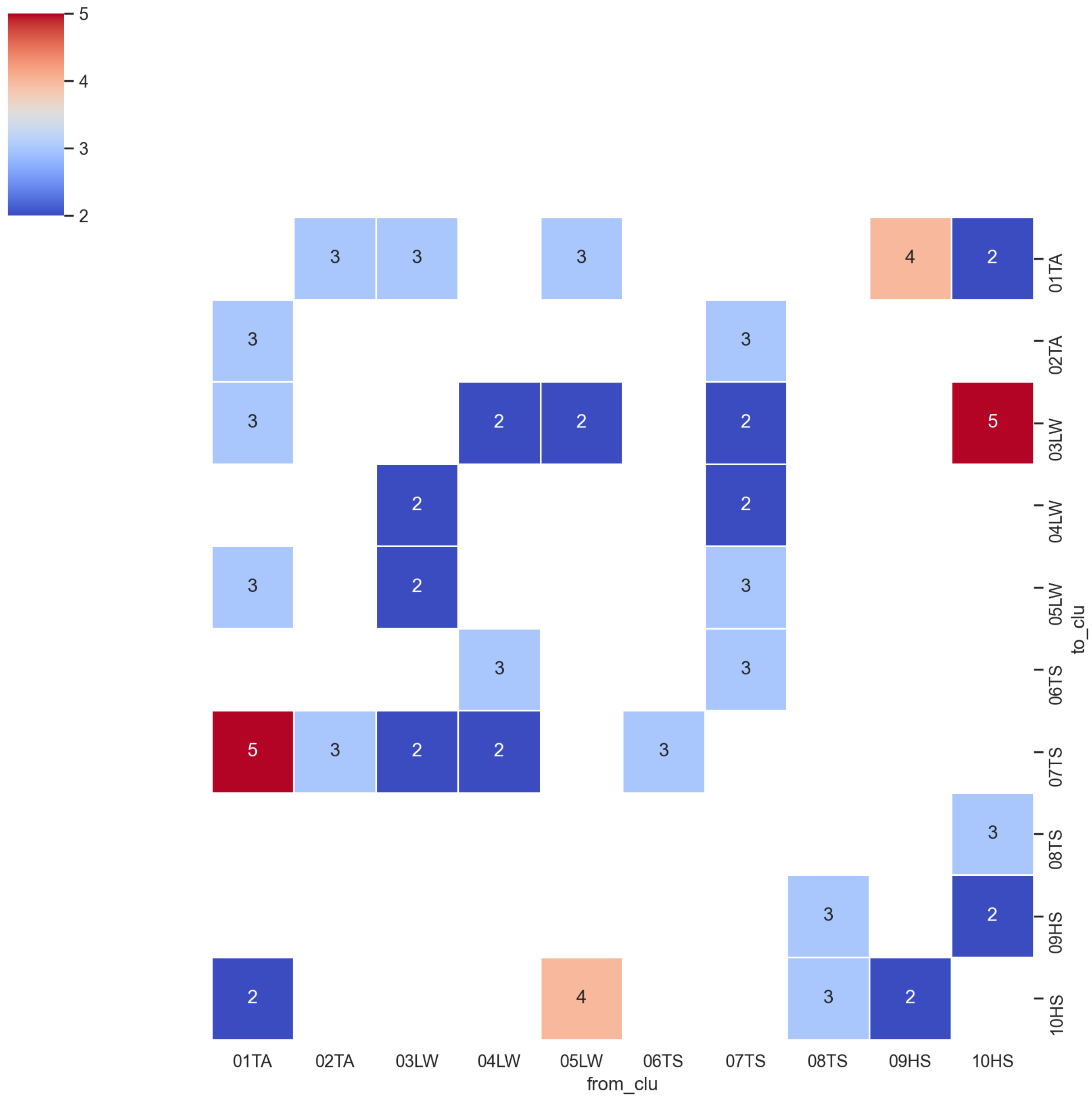


**Figure S6. Pruned Distance from Umap Projection 3d** Euklidian distance of the 3d Projection of the temporal profiles of Eukaryotes based on the cluster centroids. In contrast to Figure S5, only edge values of significant normalized mutual information are shown.

### 2.3 Pearson Correlation and normalized mutual information validated on Paramecium and Didinium

To validate our modification of the CCM using normalized mutual information instead of Pearson correlation, we use a reference dataset of abundance time series of *Paramecium aurelia* and *Didinium nasutum* as reported in (Veilleux, 1979).

The results shown in Figure S7 demonstrate bidirectional coupling between Didinium and Paramecium, consistent with established principles. Furthermore, the marked increase in accuracy, as measured by the normalized mutual information (t-test, $p < 0.05$), when mapping Didinium from the Paramecium time series compared to the reverse direction (Figure S7 B,C), implies that the predator's abundance patterns have a more significant impact on the prey's abundance than vice versa (Sugihara et al., 2012). This statement aligns with the experimental framework and the results obtained by using the unmodified method with Pearson correlation in our experimental setup, where both demonstrate a considerable correlation surpassing 0.775 ($p < 9.7 \cdot 10^{-12}$) Figure S7.

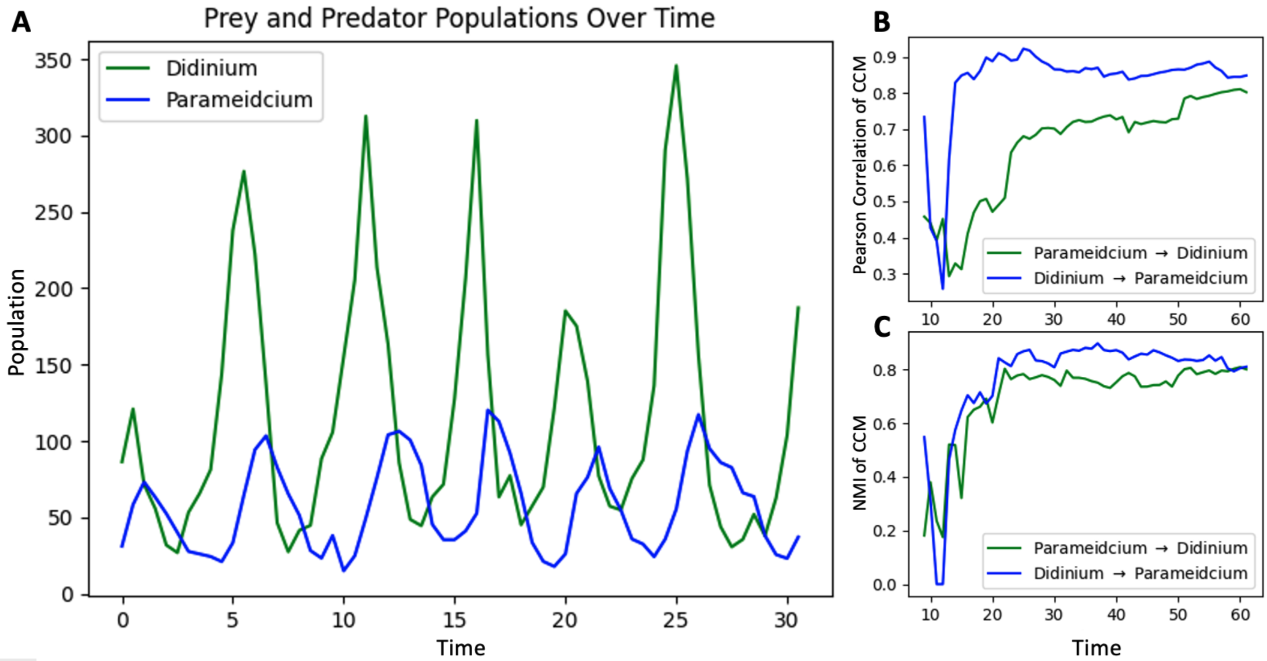


**Figure S7. Pearson Correlation and normalized mutual information CCM shows Asymmetrical bidirectional coupling for experimental setup A:** Abundance of Paramecium and Didinium in no. ind/ml) **B:** CCM (Pearson correlation) **C:** CCM (Normalized mutual information) of Paramecium and Didinium with increasing time-series length L=62. The pattern suggests top-down predator control.

### 2.4 Co-Occurence benefits Causation for our data

**Table S1. Overview CCM Variants Random, CON and Non CON.**

| cols | mean | median | std | sum | count |
|---|---|---|---|---|---|
| random | 0.171583 | 0.166083 | 0.163548 | 2960.488049 | 17254 |
| con_ccm | 0.304431 | 0.315595 | 0.203863 | 5252.660375 | 17254 |
| non_con_ccm | 0.191099 | 0.202134 | 0.165500 | 3297.223995 | 17254 |

**Table S2. Overview CCM Variants Random, CON and Non CON.**

| cols | test | alternative | test_stat | p-value | significants stars |
|---|---|---|---|---|---|
| random vs. con_ccm | Kolmogorov-Smirnov | two-sided | 0.278370 | 0.000000e+00 | *** |
| random vs. non_con_ccm | Kolmogorov-Smirnov | two-sided | 0.059349 | 0.000000e+00 | *** |
| non_con_ccm vs. con_ccm | Kolmogorov-Smirnov | two-sided | 0.247885 | 0.000000e+00 | *** |

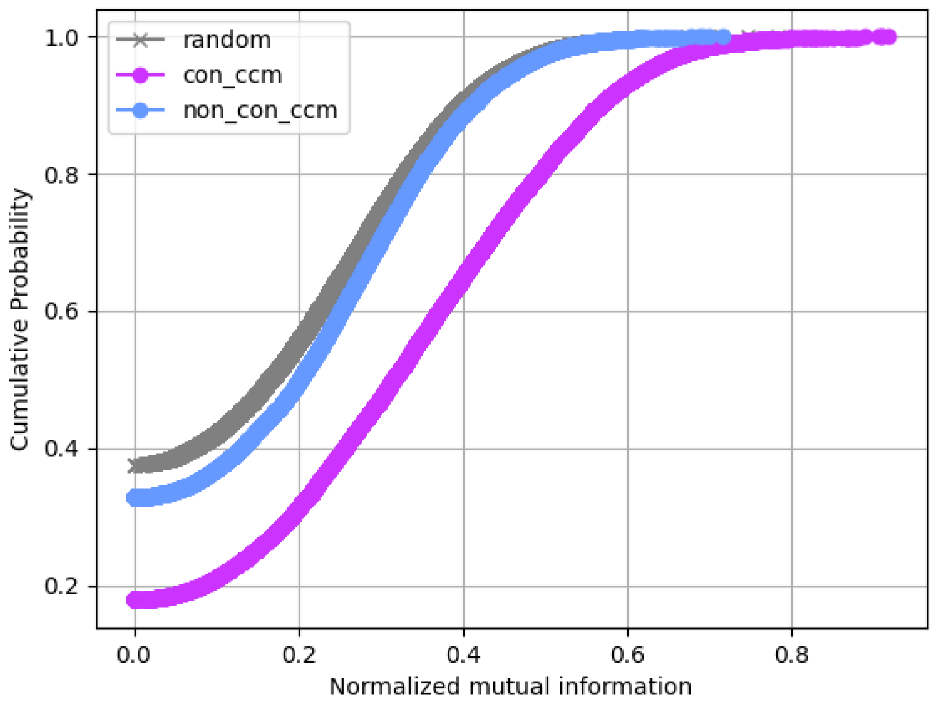


**Figure S8. Co-Occurrence benefits Interactions - CDF** We create Cumulative Distribution Function (CDF) plots for the normalized mutual information of the following three set-ups: (i): Random Matrix serving as the Null hypothesis, (ii): The intersection of the Co-Occurrence network (CON) and the Convergent Cross Mapping Network (CCMN) and (iii) the set difference of the CCMN and CON.

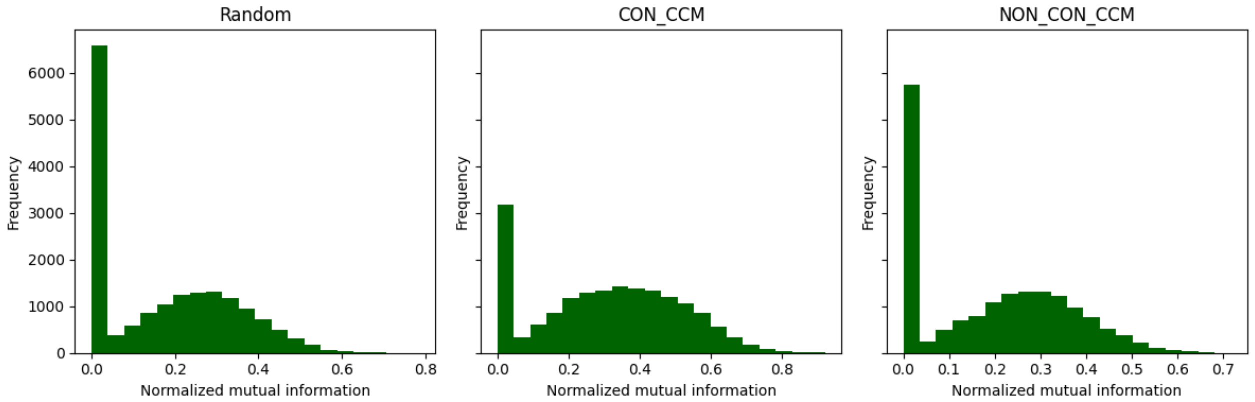


**Figure S9. Co-Occurrence benefits Interactions - Histogram** We create Frequency plots (Histograms) for the normalized mutual information of the following three set-ups: (i): Random Matrix serving as the Null hypothesis, (ii): The intersection of the Co-Occurrence network (CON) and the Convergent Cross Mapping Network (CCMN) and (iii) the set difference of the CCMN and CON.

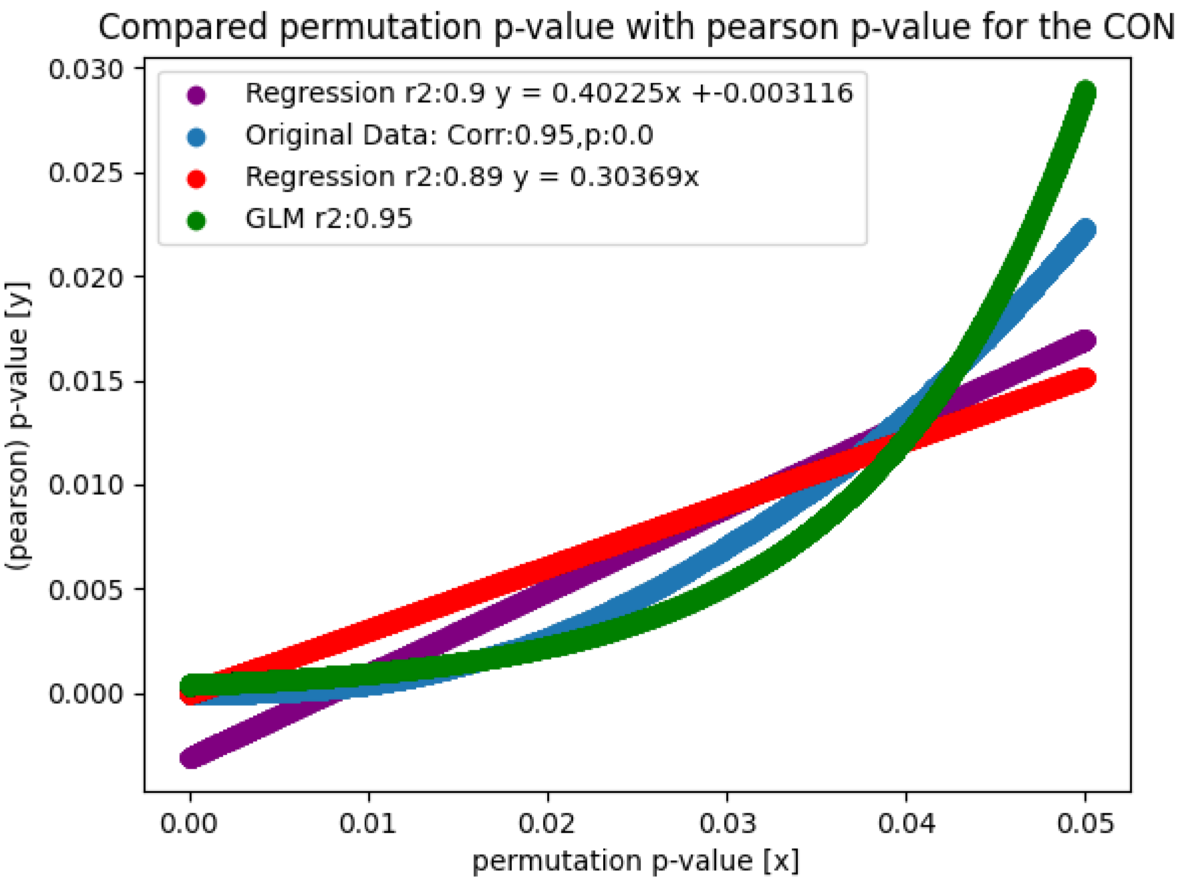


**Figure S10. Method validation of the Ψ p-value implementation.** We compared the Ψ p-value with Pearson p-value on the CON (p>0.05, corr>=0). The original data are shown in blue, the linear regression model with bias in purple and without in red. The green line present a generalized linear model of the poisson family.

### 2.5 Centrality Closeness and normalized mutual information for stable ASVs

### 2.6 Method Validation: P-value

We tested the proposed non-parametric p-value by comparing the p-value with the p-value obtained from Pearson correlation for the CON (p$<$0.05, corr $>$ 0). Our implementation was highly correlated with the p-values obtained from the Pearson correlation (corr: 0.95, p-value 0). In addition we calculated two linear regression models one with bias and one with out ($R^2$: 0.9, 0.89) and a Generalized Linear Model (GLM) of the Poisson family ($R^2$: 0.95).

### 2.7 Method Validation: CCM with NMI

To validate our modification of the CCM using normalized mutual information instead of pearson correlation, we use a reference dataset of abundance time series of *Paramecium aurelia* and *Didinium nasutum* as reported in (Veilleux, 1979). The results shown in Figure S7demonstrate bidirectional coupling, consistent with established principles. Moreover, the significant higher accuracy measured by the normalized mutual information (t-test, $p < 0.05$) in mapping Didinium from the Paramecium time series as opposed to the opposite direction (Figure S7 B,C) suggests that top-down control by the predator, Didinium, is more powerful than bottom-up control by the prey, Paramecium (Sugihara et al., 2012). This statement aligns with the experimental framework and the results obtained by using the unmodified method with Pearson correlation in our experimental setup, where both demonstrate a considerable correlation surpassing 0.775 ($p < 9.7e^{-12}$) (Figure S7).

### 2.8 Winter reset

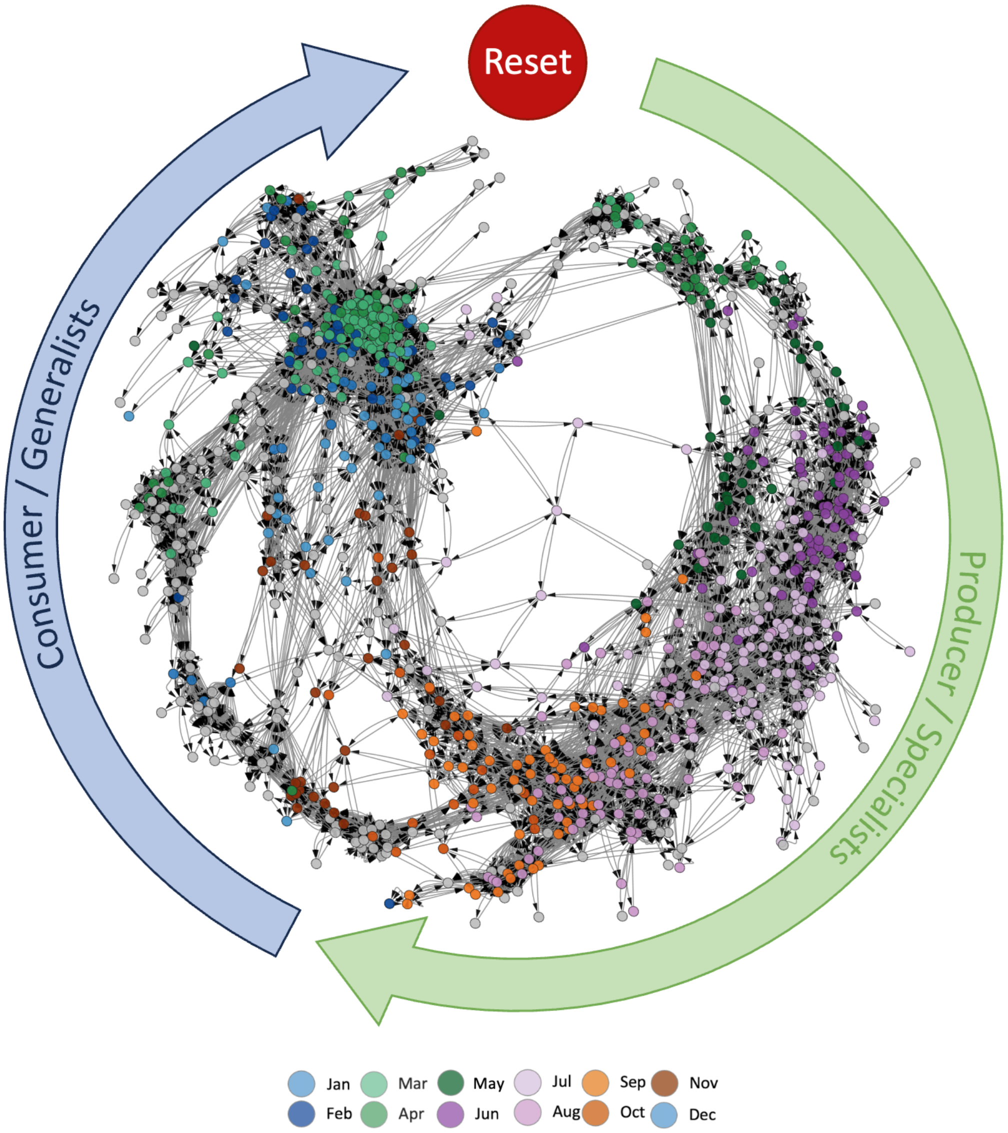


**Figure S11. Unveiling ecological dynamics: Cross Convergence Mapping Network projected Co-occurrence Network reveals causal relationships among ASVs over a 4-year period, featuring producers, consumers, and Winter Reset Events.** Cross Convergence Mapping Network Projected co-occurrence network. This network diagram illustrates the causal relationships between ASVs (amplicon sequence variants) over a 4-year observation period. Nodes represent ASVs, with arrowed edges indicating the direction of causal influence from source to target nodes. Edge weights are determined by normalised mutual information. Grey nodes indicate ASVs with no significant causal influence ($p < 0.05$). Node colours indicate the season of maximum abundance, while large arrows framing the network distinguish periods of producers (specialists) or consumers (generalists). The presence of a red reset button indicates the winter reset event.

### 3 ENERGY LANDSCAPE ANALYSIS

#### Projection of different Environmental Traits onto Phytoplankton Communities

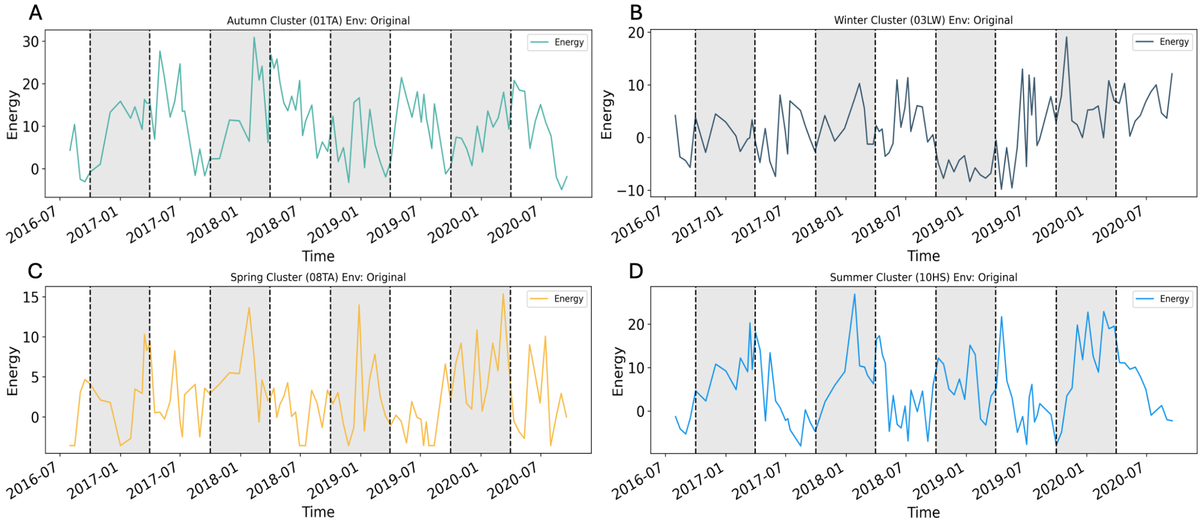


**Figure S12. Energy landscape analysis illustrating seasonal dynamics of community energy levels over a four-year span** (08.2016 - 09.2020), with each subplot depicting a distinct season (autumn, winter, spring, summer). The energy curves, colored to represent specific clusters, are based on abundance and environmental data, as indicated. The x-axis denotes time, encompassing the specified period, while environmental data integration underscores the complexity of ecological interactions shaping energy distributions.

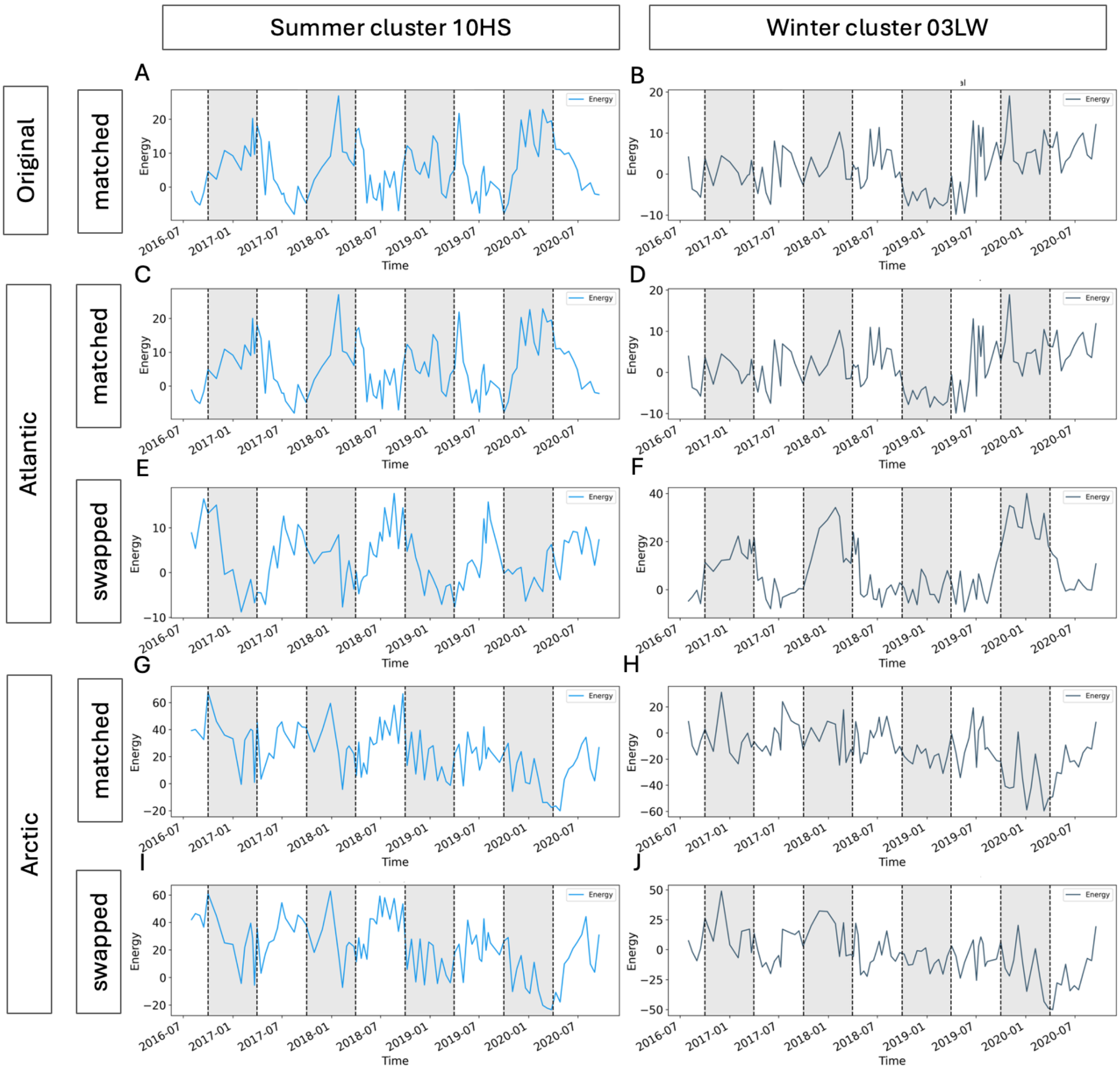


**Figure S13. Seasonal variations in community energy levels depicted through energy landscape analysis over a four-year period** (08.2016 - 09.2020), with distinct subplots representing winter and summer seasons. Each energy curve, colored to denote specific clusters, integrates abundance and environmental data, as indicated. Projections utilize extreme arctic winter and summer environmental data, applied to corresponding clusters (labeled as matched), alongside Atlantic median values over fitting seasons (May, July, June for summer and December, January, February for winter). Deviations between season and environmental condition are labeled as swapped.

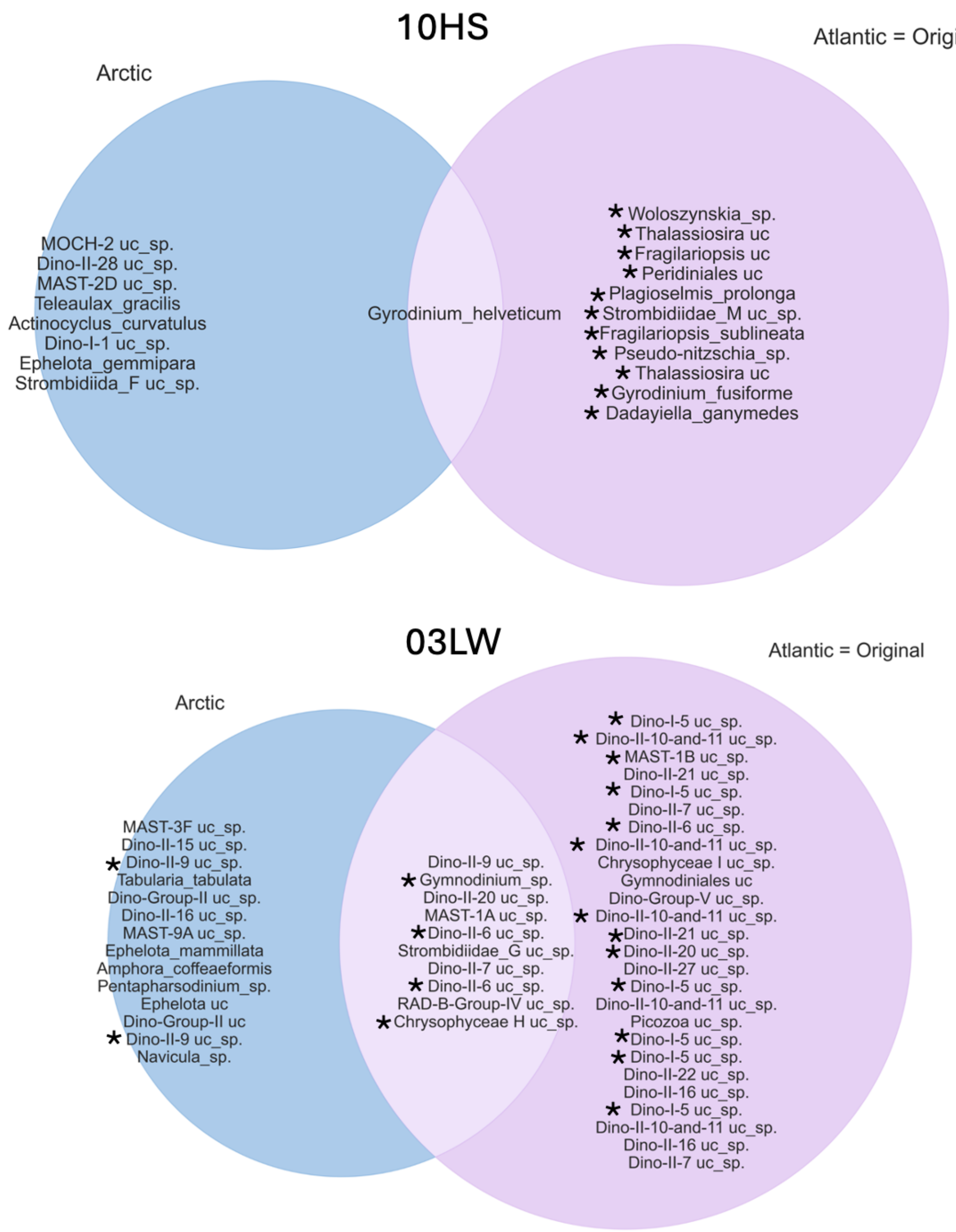


**Figure S14. Overlap between predicted potential stable species for the Atlantic projection, Arctic projection, and original data**. **A**: Venn diagram illustrating the overlap of predicted potential stable species for the winter cluster 03LW. **B**: Venn diagram illustrating the overlap of predicted potential stable species for the summer cluster 10HS. The stars indicates that these stable ASVs are also potential keystone species.

**Table S3. (Atlantic) ASV identified as a potential keystone species for clusters 10HS, 03LW projected to the Atlantic environmental parameters.** The taxa, clusters, raw abundance, proportion of total raw abundance and cluster abundance, and proportion of cluster raw abundance are shown aggregated over the 4-year observation period. The column, significance, indicates if this ASV (Nodes) has at least one significant CCM connection measured in normalized mutual information.

| Nodes | Phylum | Class | Genus | Species | Cluster | rel. Abundance | Closeness Centrality |
|---|---|---|---|---|---|---|---|
| euk_asv19 | Ochrophyta | Bacillariophyta | Thalassiosira | *Thalassiosira uc* | 10HS | 0.010 | 0.463 |
| euk_asv12 | Ochrophyta | Bacillariophyta | Fragilariopsis | *Fragilariopsis uc* | 10HS | 0.009 | 0.521 |
| euk_asv29 | Ciliophora | Spirotrichea | Strombidiidae_M uc | *Strombidiidae_M uc_sp.* | 10HS | 0.007 | 0.499 |
| euk_asv35 | Ochrophyta | Bacillariophyta | Fragilariopsis | *Fragilariopsis_sublineata* | 10HS | 0.007 | 0.533 |
| euk_asv28 | Ochrophyta | Bacillariophyta | Thalassiosira | *Thalassiosira uc* | 10HS | 0.006 | 0.473 |
| euk_asv54 | Dinoflagellata | Dinophyceae | Gyrodinium | *Gyrodinium_fusiforme* | 10HS | 0.006 | 0.469 |
| euk_asv24 | Ochrophyta | Bacillariophyta | Pseudo-nitzschia | *Pseudo-nitzschia_sp.* | 10HS | 0.006 | 0.446 |
| euk_asv60 | Dinoflagellata | Dinophyceae | Woloszynskia | *Woloszynskia_sp.* | 10HS | 0.005 | 0.489 |
| euk_asv73 | Cryptophyta | Cryptophyceae | Plagioselmis | *Plagioselmis_prolonga* | 10HS | 0.005 | 0.513 |
| euk_asv52 | Dinoflagellata | Dinophyceae | Peridiniales uc | *Peridiniales uc* | 10HS | 0.004 | 0.486 |
| euk_asv125 | Ciliophora | Spirotrichea | Dadayiella | *Dadayiella_ganymedes* | 10HS | 0.003 | 0.504 |
| euk_asv87 | Radiolaria | RAD-B | RAD-B-Group-IV uc | *RAD-B-Group-IV uc_sp.* | 03LW | 0.002 | 0.489 |
| euk_asv79 | Ochrophyta | Chrysophyceae | Chrysophyceae H uc | *Chrysophyceae H uc_sp.* | 03LW | 0.002 | 0.409 |
| euk_asv236 | Dinoflagellata | Syndiniales | Dino-II-9 uc | *Dino-II-9 uc_sp.* | 03LW | 0.001 | 0.438 |
| euk_asv213 | Dinoflagellata | Dinophyceae | Gymnodinium | *Gymnodinium_sp.* | 03LW | 0.001 | 0.434 |
| euk_asv198 | Dinoflagellata | Syndiniales | Dino-II-6 uc | *Dino-II-6 uc_sp.* | 03LW | 0.001 | 0.433 |
| euk_asv411 | Dinoflagellata | Syndiniales | Dino-II-20 uc | *Dino-II-20 uc_sp.* | 03LW | 0.001 | 0.428 |
| euk_asv615 | Dinoflagellata | Syndiniales | Dino-II-20 uc | *Dino-II-20 uc_sp.* | 03LW | 0.001 | 0.508 |
| euk_asv511 | Dinoflagellata | Syndiniales | Dino-II-10-and-11 uc | *Dino-II-10-and-11 uc_sp.* | 03LW | 0.001 | 0.400 |
| euk_asv553 | Dinoflagellata | Syndiniales | Dino-II-21 uc | *Dino-II-21 uc_sp.* | 03LW | 0.001 | 0.430 |
| euk_asv780 | Dinoflagellata | Syndiniales | Dino-I-5 uc | *Dino-I-5 uc_sp.* | 03LW | 0.000 | 0.410 |
| euk_asv1293 | Dinoflagellata | Syndiniales | Dino-II-10-and-11 uc | *Dino-II-10-and-11 uc_sp.* | 03LW | 0.000 | 0.407 |

**Table S4. (Simulated Arctic conditions) ASV identified as potential keystone species for cluster 10HS, 03LW projected to the arctic environmental parameters.** The taxa, cluster, raw abundance, proportion of total raw abundance and cluster abundance, and the proportion of cluster raw abundance are shown aggregated over the 4-year observation period. The column, significance, indicates if this ASV (Nodes) has at least on significant CCM connection measured in normalized mutual information.

| Nodes | Phylum | Class | Genus | Species | Cluster | rel. Abundance | Closeness Centrality |
|---|---|---|---|---|---|---|---|
| euk_asv75 | Dinoflagellata | Syndiniales | Dino-I-1 uc | *Dino-I-1 uc_sp.* | 10HS | 0.004 | 0.435 |
| euk_asv161 | Cryptophyta | Cryptophyceae | Teleaulax | *Teleaulax_gracilis* | 10HS | 0.002 | 0.457 |
| euk_asv223 | Pseudofungi | MAST-2 | MAST-2D uc | *MAST-2D uc_sp.* | 10HS | 0.002 | 0.418 |
| euk_asv579 | Dinoflagellata | Syndiniales | Dino-II-28 uc | *Dino-II-28 uc_sp.* | 10HS | 0.001 | 0.420 |
| euk_asv87 | Radiolaria | RAD-B | RAD-B-Group-IV uc | *RAD-B-Group-IV uc_sp.* | 03LW | 0.002 | 0.489 |
| euk_asv79 | Ochrophyta | Chrysophyceae | Chrysophyceae H uc | *Chrysophyceae H uc_sp.* | 03LW | 0.002 | 0.409 |
| euk_asv236 | Dinoflagellata | Syndiniales | Dino-II-9 uc | *Dino-II-9 uc_sp.* | 03LW | 0.001 | 0.438 |
| euk_asv213 | Dinoflagellata | Dinophyceae | Gymnodinium | *Gymnodinium_sp.* | 03LW | 0.001 | 0.434 |
| euk_asv291 | Dinoflagellata | Syndiniales | Dino-Group-II uc | *Dino-Group-II uc_sp.* | 03LW | 0.001 | 0.396 |
| euk_asv615 | Dinoflagellata | Syndiniales | Dino-II-20 uc | *Dino-II-20 uc_sp.* | 03LW | 0.001 | 0.508 |
| euk_asv803 | Opalozoa | MAST-3 | MAST-3F uc | *MAST-3F uc_sp.* | 03LW | 0.001 | 0.407 |
| euk_asv602 | Dinoflagellata | Syndiniales | Dino-II-9 uc | *Dino-II-9 uc_sp.* | 03LW | 0.000 | 0.454 |
| euk_asv1088 | Dinoflagellata | Syndiniales | Dino-II-16 uc | *Dino-II-16 uc_sp.* | 03LW | 0.000 | 0.428 |
| euk_asv837 | Dinoflagellata | Syndiniales | Dino-Group-II uc | *Dino-Group-II uc* | 03LW | 0.000 | 0.502 |

### 3.1 Connections of Stable ASVs in the CCM Network

We filtered all edges connecting two ASVs nodes, where at least one node is a stable ASV. Next, we determined the number of edges in its cluster and other clusters. Then, we calculated the normalized mutual information from stable ASVs to ASVs and from ASVs of their cluster. We compared the stable ASV from a winter cluster example, 03LW, and a summer cluster example, 10HS, under Atlantic and Arctic environmental conditions.

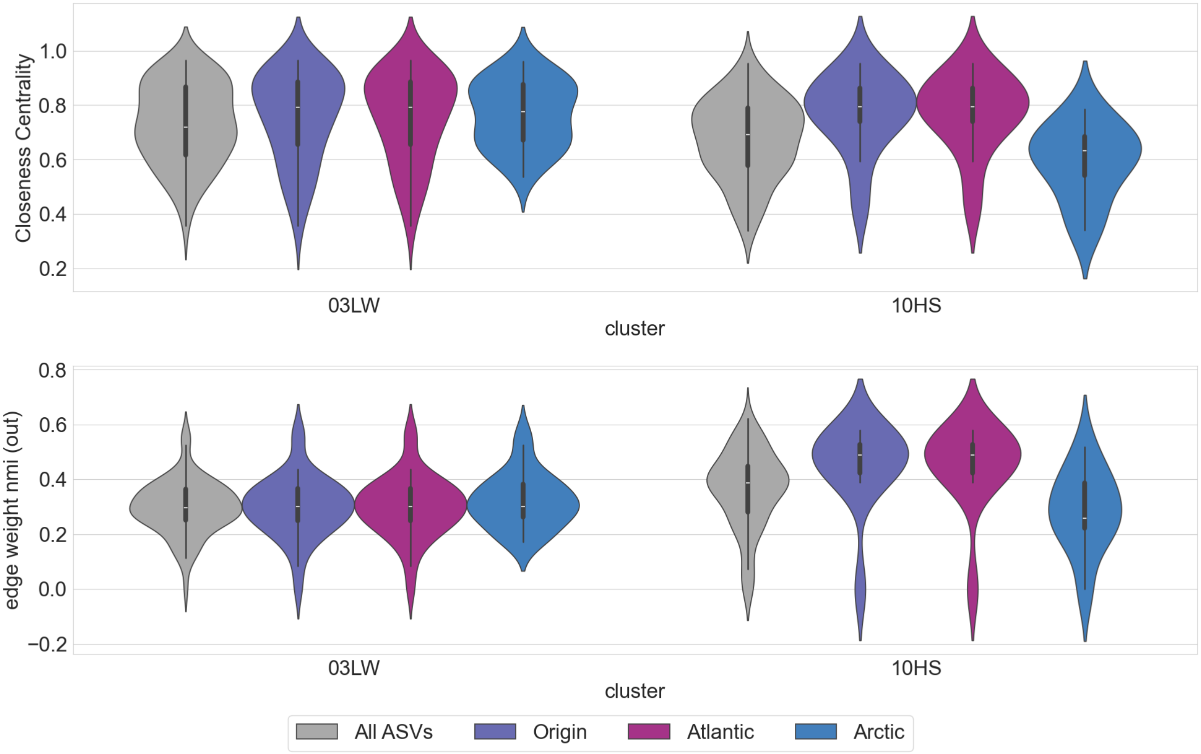


**Figure S15. Closeness Centrality and edge weight for All ASVs, Origin, Arctic and Atlantic Environments stable state ASVs. A):** The Closeness Centrality for stable ASVs for Atlantic Environmental condition vs non stable state ASVs. The stable ASVs from the arctic environment are indicated with blue, the stable ASVs from the atlantic environment are purple and the non stable ASVs are black. **Comparison of connection and interaction for Arctic and Atlantic winter and summer. B):** The mean over normalized mutual information to the stable ASVs from other ASVs.
